# Evolutionary new centromeres in the snowy owl genome putatively seeded from a transposable element

**DOI:** 10.1101/2024.07.05.602039

**Authors:** H. T. Baalsrud, B. Garmann-Aarhus, E. L. G. Enevoldsen, A.K. Krabberød, D. Fischer, A. Tooming-Klunderud, M. Skage, M. Árnyasi, S. R. Sandve, K.S. Jakobsen, R. Nielsen, S. Boessenkool, O. K. Tørresen

## Abstract

Comparative genomic studies in birds have revealed that bird genomes are relatively repeat-poor and stable in terms of karyotype, size, and gene synteny/collinearity compared to other vertebrates. One notable exception is the owls, with cytogenetic studies demonstrating large variations in karyotypes and the evolution of unusual centromeric satellite repeats in some species. However, there has so far not been an investigation into genome architecture and repeat landscape of owls. Here, we present a chromosome-level genome assembly for the snowy owl (*Bubo scandiacus*). We find that the repeat DNA content in the relatively large snowy owl genome (1.6 Gb) is among the highest reported for any bird genome to date (28.34% compared to an average of ∼10% in other birds). The bulk of the snowy owl genomic repeat landscape consists of centromeric satellite DNA, which appears to have originated from an endogenous retrovirus (ERV1). Using gene collinearity analyses we show that the position of these evolutionary new centromeres (ECNs) are not homologous with chicken centromeres, and are located in regions with collinearity breaks to other bird genomes due to chromosomal rearrangements. Our results support rapid transposable element-driven evolution of lineage-specific centromeres, which could have played a role in reproductive isolation and speciation of the snowy owl.

## Introduction

The recent rapid accumulation of high-quality genome assemblies across the Tree of Life has significantly advanced our understanding of genome evolution. However, even within well-studied taxonomic groups like birds (Jarvis et al. 2014; Zhang et al. 2014; Feng et al. 2020; Stiller et al. 2024), there are still lineages that lack high-quality long-read reference genomes and thereby represent gaps in our understanding of chromosomal architecture and genomic repeats (Braun et al. 2019; Feng et al. 2020). The owls (Strigiformes) are one such understudied group.

Earlier cytogenetic studies of the owls have demonstrated fascinating aspects of chromosome evolution within both of its two families, the barn owls (Tytonidae) and the true owls (Strigidae). Tytonidae seem to lack the classic avian genome organization of micro-and macro chromosomes (Rebholz et al. 1993). Strigidae are also unusual by having large genome sizes compared to other birds (average 1.6 Gb), with several species displaying large heterochromatin blocks in the centromeric regions of their chromosomes (Yamada et al. 2004). These regions have been shown to harbor satellite repeats unique to the true owls, which cluster phylogenetically by species (Yamada et al. 2004). Centromeric regions are known hotspots for chromosomal rearrangements in other systems (Smalec et al. 2019; Ola et al. 2020). In owls, however, the genomic impact of these centromeric repeats is still lacking.

Here we present a chromosome-level reference genome assembly for the iconic snowy owl (*Bubo scandiacus*) as part of the Earth Biogenome Project Norway initiative (https://www.ebpnor.org/english/). We have used a combination of chromatin interaction mapping (Hi-C) and long reads from both Pacific Biosciences (PacBio) and Oxford Nanopore Technologies (ONT), which allows us to characterize the genomic repeat landscape of this species. By comparing the snowy owl assembly with representative chromosome-level assemblies from other birds, we investigate the following three questions: What is the overall repeat content in the snowy owl genome compared to other birds? How are the centromeric satellite DNA and other classes of repeats distributed across the snowy owl repeat landscape? And what is the impact of the unique centromere-associated repeat landscape on synteny to other avian genomes? This in-depth genomic investigation of the repeat landscape of the snowy owl will lift our knowledge of bird genome evolution. Furthermore, this study can facilitate future research and management on the snowy owl (*Bubo scandiacus*), which is a key apex predator and an indicator species for the health of the Arctic ecosystems.

## Results

### A chromosome-level genome assembly for the snowy owl

We obtained a highly contiguous, phased, diploid snowy owl assembly (Figure 1) by leveraging the high accuracy of PacBio HiFi reads, the length of ONT reads, and linkage information in Hi-C reads (Figure S1). While the expectation is that the assembly is mostly phased within each chromosome, the chromosomes themselves are not phased with respect to each other, that is, chromosomes one and two in haplotype one might not be from the same parent. Recognizing that the two assemblies are technically pseudo-haplotypes if defining haplotypes as unions of segments of DNA inherited from the same parent, we refer to them as haplotypes (hap1 and hap2). The resulting assembly was 1.6Gb and had a contig N50 of 16.5 Mb and scaffold N50 of 51 Mb for hap1 (Figure 1). Both sex chromosomes were assigned to hap1 for convenience. Completeness based on k-mers was 96.5 % for hap1, 86.4 % for hap2 and 98.4 % for both combined. Hap 1 had 97.3 % complete genes based on BUSCO, while hap2 had 90.1 % BUSCO completeness. Using a manual curation process, which included homology comparisons to the assemblies of chicken and zebra finch (see Methods for details), we identified 32 and 30 autosomes in hap1 and hap2, respectively. Cytogenetic studies have previously found that the chromosome number is 2N=82 in the snowy owl (Sasaki et al. 1994), which indicates that 10 pairs of the microchromosomes were not distinguished from unplaced scaffolds in our assembly. This is typical for vertebrate species with microchromosomes, which tend to cluster together during meiosis and are notoriously difficult to characterize (Waters et al. 2021; Huang et al. 2023).

**Figure 1.**
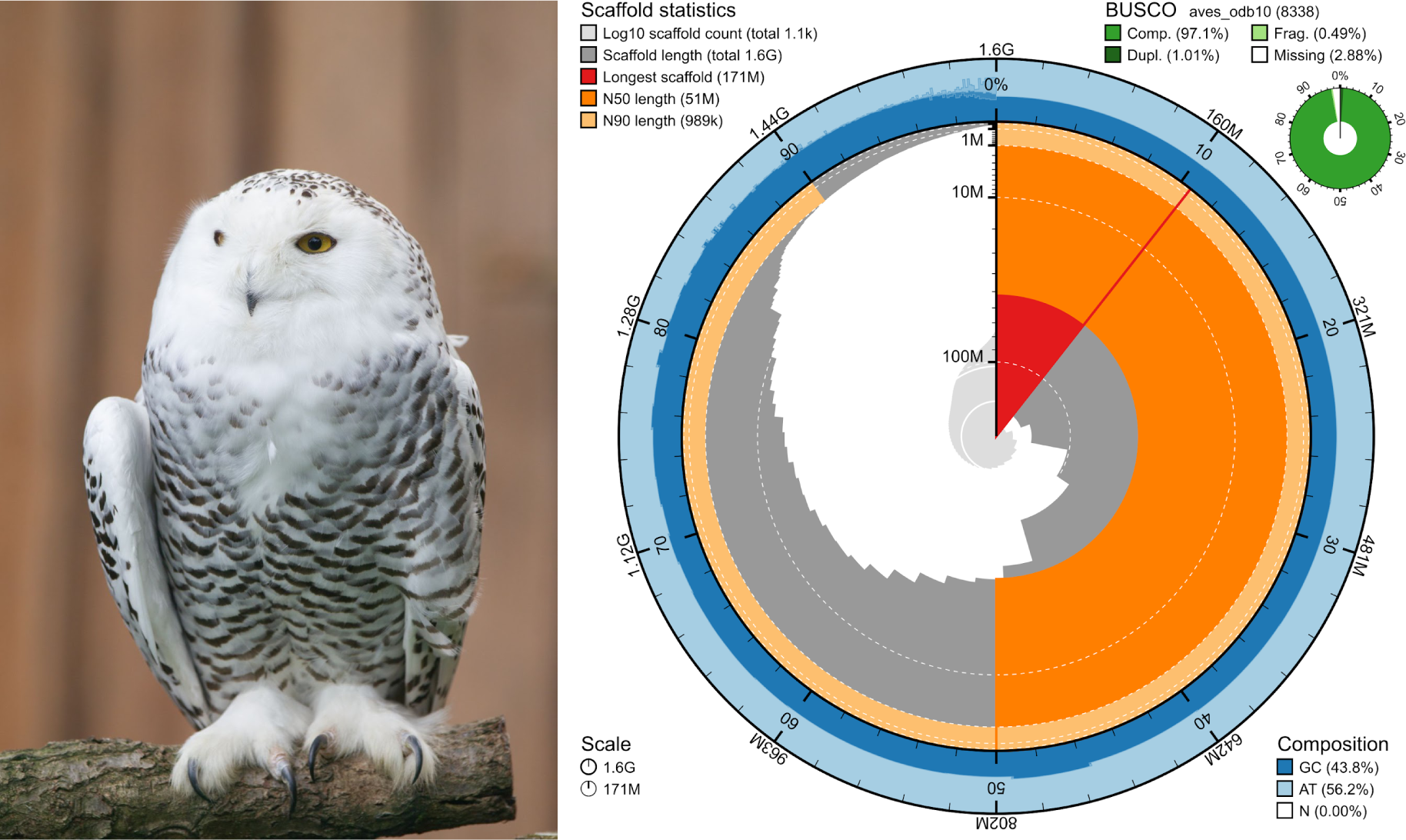
Genome assembly of *Bubo scandiacus, bBubSca1.1: metrics.* The BlobToolKit Snailplot shows a graphical representation of the assembly metrics for hap1 presented in Table S1. The snailplot circle represents the assembly, with the red line represents the largest scaffold and all other scaffolds arranged clockwise subsequent to this, with scaffold N50 and N90 shown according to legend. The blue areas show base composition and the smaller green circle shows BUSCO gene completeness. The picture of the snowy owl female sequenced in this project was taken by Wolfgang Holtmeier at the Raptor Center & Wildlife Zoo, Hellenthal.

Our comparisons between hap1 and hap2 revealed several putative inversions, duplications, and translocations (Figure S2). In total, we found 159 syntenic regions (∼1 Gb), 16 inversions (∼45 Mb), 155 translocations (∼48 Mb), 121 duplications in hap1 (∼1 Mb), and 194 duplications in hap2 (∼2 Mb). The largest inversions were found on chromosomes 2, 3, 4, and 5.

### Phylogenetic analyses based on orthologs

In order to study genome evolution in the snowy owl, we selected five other species with high-quality assemblies in the Telluraves lineage for comparison: downy woodpecker (*Dryobates pubescens*), Northern Carmine bee-eater (*Merops nubicus*), Northern goshawk (*Accipiter gentilis*), California condor (*Gymnogyps californianus*), and barn owl (*Tyto alba*), as well as two outgroup species: chicken (*Gallus gallus*) and zebra finch (*Teaniopygia guttata*). Our phylogenetic tree placed Strigiformes as a sister clade to Accipitriformes (Figure 2 and Figure S7), which is concordant with the topology most recently published phylogenetic tree for birds (Stiller et al. 2024).

**Figure 2.**
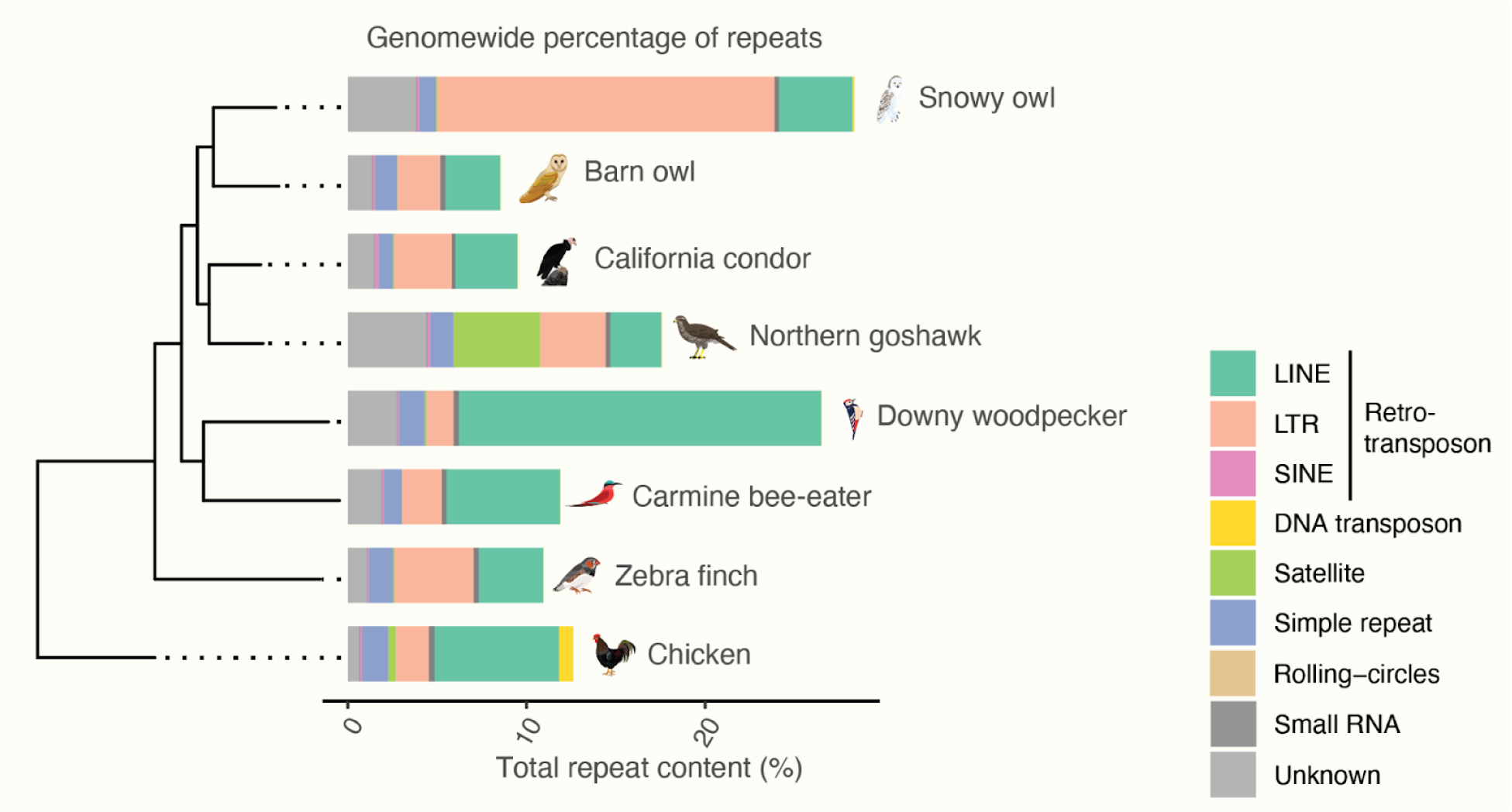
Repeat content of a selection of avian genome assemblies. Repeat content for all species, with different classes colored according to legend. The phylogenetic tree was generated using Orthofinder (See Data Availability). The figure was made in R and modified in Illustrator.

### The snowy owl repeat landscape

*De novo* repeat modeling and detection revealed that 28.34 % of the snowy owl genome consists of repeat DNA (Figure 2), which is significantly higher than the average reported repeat content of ∼10% across birds (Kapusta and Suh 2017). This makes the snowy owl genome a notable outlier among birds in company with the downy woodpecker (Kapusta and Suh 2017; Manthey et al. 2018) and Bell’s sparrow (*Artemisiospiza belli*) (Benham et al. 2023). Unlike the two latter species, the size of the snowy owl genome is significantly larger than the average avian genome size (1.6 Gb vs an average of ∼1 Gb), suggesting that repeat expansions in the snowy owl are driving genome size evolution. To explore this further, we annotated repeat DNA across all Telluraves and outgroup species in our study and found that the snowy owl had the highest absolute repeat content (Figure 2, Supplementary Table X). The type of repeat DNA also differed extensively between snowy owl and the other species. In terms of base pairs, most of the repeats in the snowy owl genome were annotated as LTR retrotransposons (LTRs) (Figure 2 and Figure S3). A staggering 255 Mb of the LTRs consist of one type; endogenous retrovirus 1 (ERV1), class family-0. In contrast, in the other bird species with high repeat content, the downy woodpecker and Bell’s sparrow, there has been an expansion of CR1 LINE elements (Kapusta and Suh 2017; Benham et al. 2023). The barn owl, California condor, and Northern goshawk genomes do not appear to have similar expansions of LTRs (Figure 2), which seems to be unique to the snowy owl genome. The Northern goshawk genome is also quite repeat-rich (17.53%) by avian standards, with a large proportion of satellite DNA.

### Evolutionary new centromeres in the snowy owl genome

To understand the significance of the extensive expansion of ERV1 Class family-0 in the snowy owl genome we investigated this family in more detail. We compared the family-0 consensus sequence (length = 305bp) to the NCBI nucleotide database using BLAST and found 74 highly significant hits (e-value > 1e27, percentage identity > 83.24) against centromeric repeat region from six species within Strigidae (including snowy owl) (Yamada et al. 2004), and no hits against any other species. This suggests the highly abundant family-0 repeats found in our genome assembly are the centromeric satellite DNA identified in (Yamada et al. 2004), and moreover, that these centromeric repeats are evolutionary new centromeres (ENC) unique to the Strigidae family. Yamada et al. (2004) used probes designed with cloned sequences from another species (*Bubo blakistoni*) and estimated that 9.6% of the snowy owl genome consists of (15.8% of hap1).

To investigate the physical location of centromeric repeats in the snowy owl genome, we estimated the density of different classes of repeats in 200 kb windows, dividing the LTR class into two groups, family-0 and LTR-other. The family-0 repeats are not evenly spread across the genome but form large, contiguous blocks, consistent with the approximate position of centromeres found in (Yamada et al. 2004) (Figure 3). Cross-referencing the cytogenetic images in (Yamada et al. 2004) with the presence of family-0 repeats suggests that chromosomes 3, 4, and 5 are metacentric/submetacentric. The remaining chromosomes have family-0 repeats at the end and appear to be acrocentric. The two medium-sized chromosomes with a very weak fluorescence signal for the centromeric satellite DNA (Yamada et al. 2004) may correspond to chromosomes 6 and 8 (Figure 3), which have very small regions of family-0 repeats. Some of the smaller chromosomes (16, 17, 19, and 29) have no family-0 sequence and are difficult to match to the cytogenetically identified chromosomes. The absence of family-0 sequences in these smaller chromosomes may be due to misassembly, as there is 206 Mb of family-0 in unplaced scaffolds of a total 254 Mbp in the assembly.

**Figure 3:**
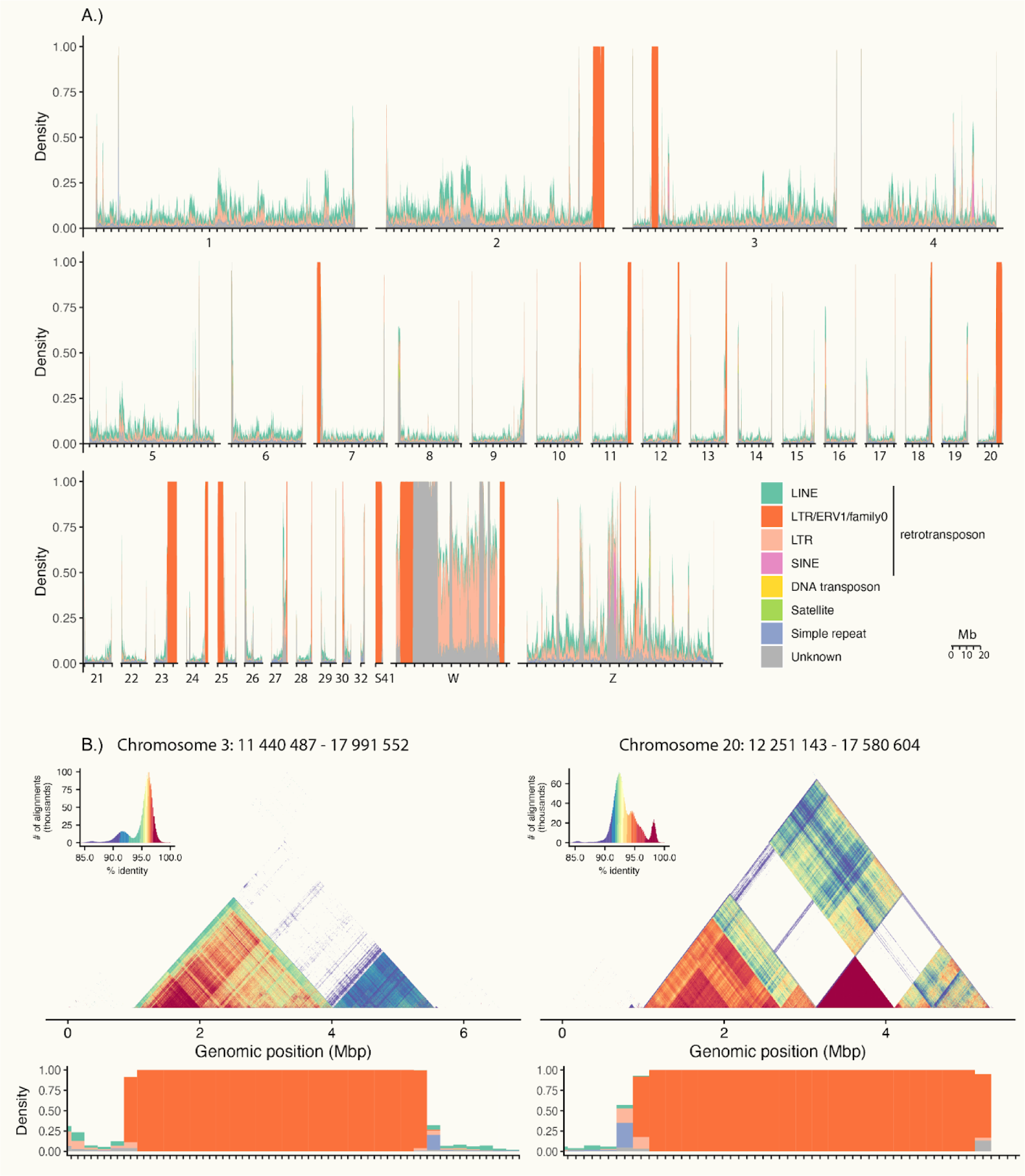
The chromosomal landscape of repeats in the snowy owl genome. A.) Densities of repeats in 200kb windows across all curated, putative chromosomes. Different repeat classes are colored according to legend. The LTR class was divided into family-0 elements and LTR other for clarity. B.) Examples of higher-order repeat structure of the putative centromere/pericentromere regions at chromosomes 3 and 20. Each region is delimited by the position of family-0 elements in addition to 1 Mb of flanking sequence on either side (except for the right region on chromosome 20 where the flanking region is 40 kbp because the chromosome ends). The heatmaps display all-by-all present identity colored according to legend, calculated for 2 kb windows using StainedGlass. The lower panel is a zoom-in of repeat density as in A.) for the same region. This plot was made in R and modified in Illustrator.

### Size and relative age distribution of snowy owl repeats

We also wanted to look at the relative timing of repeat expansions, especially the family-0 elements, in the snowy owl genome. The distribution of the percentage divergence to the consensus sequence is a good proxy for the relative age of a repeat family. We found that the distribution of percentage divergence varied both within and between repeat families in the snowy owl genome (Figure S4). In relative terms, the most recent repeat expansions seem to be SINE 5S, and also some of the simple repeats, rRNAs and LINE/L1 elements. Since SINE 5S is a non-autonomous TE it is tempting to speculate that it has piggy-backed on the LINE/L1 expansion. The ERV1s had a wide span of percentage divergence to the consensus. The ERV1-family-0 separately showed a bimodal distribution of the percentage divergence from the consensus (Figure S5B). The regions of family-0 elements ranged in size from 41 to 5 055 234 bp, with a median size of 133 006 bp (Figure S5A). This is several magnitudes larger than what would be expected for ERVs (Wells and Feschotte 2020), but is consistent with long stretches of satellite DNA. The family-0 elements form complex higher-order repeat structures in the putative centromeric regions (Figure 3B, Figure S6). In the flanking regions there is no such higher-order repeat structure, however, there are other repeats such as TEs and simple repeats (Figure 3B). The larger elements are more similar to the consensus sequence than the smaller elements (Figure S5C).

It is difficult to say whether these centromeric repeats originated by seeding of ERV1 family-0 or some other satellite sequence followed by a later invasion of ERV1 family-0 elements. Another possibility is that these centromeric satellite repeats were misannotated as ERV1s because they share sequence similarity due to convergence. We confirmed homology to ERV1 sequences using BLAST (Altschul et al. 1990), and the annotation was based on RepeatModeler2 (Flynn et al. 2020), which uses structural LTR detection and filtering methods to remove false positives and identify internal LTR regions. We did not detect any ERV genes within the family-0 elements, however, this is not unexpected in mature centromeric satellite DNA (Altemose et al. 2022; Packiaraj and Thakur 2024).

### Genomic rearrangements and syntenic collinearity

To investigate whether the ENC has had an effect on genome-wide collinearity between the snowy owl and other birds as regions surrounding centromeres are known hotspots for rearrangements (Smalec et al. 2019; Ola et al. 2020; Dobry et al. 2023), we investigated collinearity based on orthogroups using GENESPACE (Lovell et al. 2022). The overall gene collinearity including genomic regions flanking centromere sequences was generally conserved, with a few notable exceptions (Figure 4). Although collinearity was, as expected, fairly conserved between the snowy owl genome and the other avian genomes included in this study, we also found extensive genomic rearrangements between species (Figure 4). The degree of rearrangements do not necessarily only reflect evolutionary distance. For instance, there seems to have been many fissions in the barn owl genome since the split with the snowy owl, in addition to translocations indicated by the collinearity between snowy owl chr2 and barn owl scaffold 282.1. Some of these fission events may reflect assembly artifacts, as this assembly is not based on any contact map data. However, cytogenetic studies have shown that the barn owl does not have the classic division between macro- and microchromosomes and instead has a more uniform chromosome distribution. Compared to the barn owl, the snowy owl genome has more conserved collinearity with chicken and zebra finch (Figure 4).

**Figure 4:**
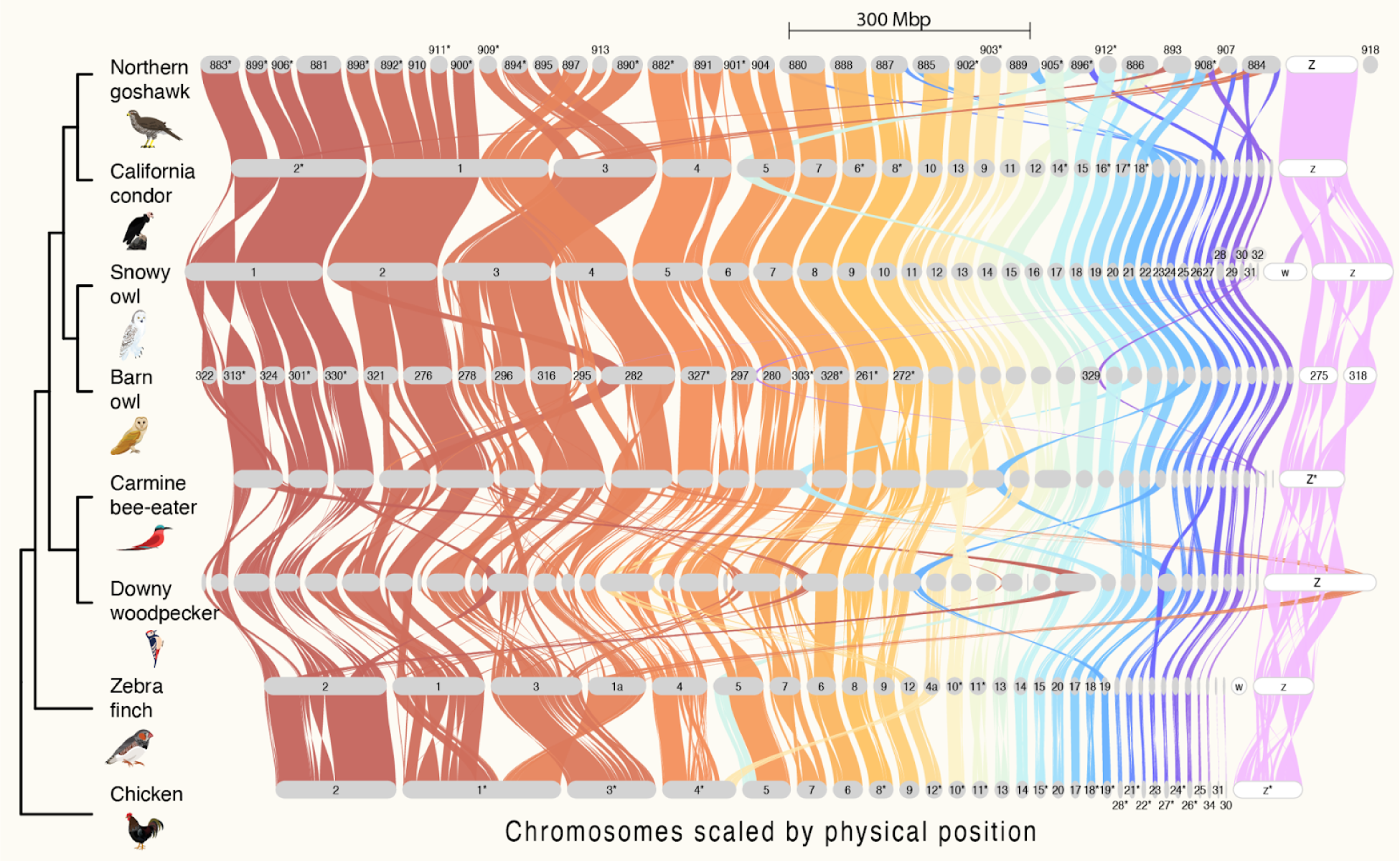
Chromosome-level syntenic network of snowy owl and representative avian assemblies. The plot was generated by GENESPACE. Genomes are ordered vertically according to phylogenetic relationships. The phylogenetic tree was generated by Orthofinder. Chromosomes are ordered horizontally to maximize collinearity with the snowy owl genome, with the sex chromosomes (Z and W) shown to the left in white. Only chromosomes or scaffolds with >100 genes and collinearity blocks > 5 genes were included in the plot. Chromosome segment sizes are scaled by Mbs according to legend. Braids illustrate gene order along the chromosome sequence. Some chromosome names were abbreviated or omitted for clarity, see supporting information for pairwise synteny comparisons with full names. *Indicate inverted chromosomes.

The snowy owl has a higher degree of collinearity with the two species belonging to Accipitriformes, especially the California condor, than with the carmine bee-eater and the downy woodpecker (Figure 4). Since the split between the snowy owl and the California condor there have been two fission events; California condor’s chromosome 1 is split into snowy owl chromosomes 2 and 4, and California condor’s chromosome 5 is split into snowy owl chromosomes 5 and 1. Northern goshawk has undergone several fissions and therefore has smaller macrochromosomes than the California condor. Part of the snowy owl W chromosome is homologous to the Z chromosome in California condor and Northern goshawk, indicating a translocation to the snowy owl W chromosome. The snowy owl W chromosome is 60 Mb, which is relatively large compared to other birds (Xu et al. 2019; Xu and Zhou 2020). Northern goshawk also has a large W chromosome (∼40Mb), whereas in the California condor W appears to be missing from the assembly even though a female was sequenced. As W is a refugium for repeats (Peona et al. 2021) it is often difficult to assemble, and collinearity comparisons of W become challenging.

The lack of large macro chromosomes also seems to characterize carmine bee-eater and downy woodpecker. In the latter species, there have been massive rearrangements, including inversions, fission/fusion events, and translocations (Figure 4). Interestingly, a large chunk of the downy woodpecker Z chromosome (∼23 Mb) is homologous to autosomes in the other birds (e.g. chicken chromosome 1). This indicates a translocation from an autosome to a sex chromosome in the ancestor of the downy woodpecker.

### Centromere position homology between the snowy owl and chicken

Centromeres are known hotspots for chromosome rearrangements (Smalec et al. 2019; Ola et al. 2020). To investigate the link between centromeres and karyotype evolution in the snowy owl, we therefore compared the snowy owl centromere positions and gene order colinearity to the well-characterized chicken genome (Huang et al. 2023). Chicken has 7 chromosomes that are submetacentric, metacentric, or sub-telocentric and all of these have large structural rearrangements in comparison to the snowy owl, including fusion/fission events and inversions (Figure 4). We then compared the chicken centromere position to collinearity breakpoints and found a complete overlap for chicken chromosomes 1, 4, 7, and 8, and near overlap for chicken chromosomes 2, 3, and 5 (Figure S8). For the snowy owl we used the position of the family-0 repeats to approximately characterize the centromere locations on the genome. The putative positions of the ENCs in the snowy owl metacentric/submetacentric chromosomes 3, 4, and 5 are not homologous with the centromere position in chicken (Figure S8). However, they do overlap with or are very close to breaks in collinearity between these two species in chromosomal regions with many rearrangements. We also investigated collinearity between the snowy owl ENCs on chromosomes 3, 4, and 5 with barn owl and found that the snowy owl centromere position overlaps with collinearity breaks, but not with fission breakpoints in the barn owl (Figure S9). There are three putative fission events in the snowy owl compared with chicken, two of which occurred in the chicken centromeric region (Chicken chromosomes 1 and 4)(Figure 4 and Figure S8). Only snowy owl chromosomes 2 and 12 evolved ENCs at the fission breakpoints resulting in acrocentric chromosomes, whereas chromosomes 4 and 5 evolved ENCs in a genomic region not homologous to the fission breakpoints.

## Discussion

We have produced a chromosome-level assembly for the snowy owl using a combination of PacBio HiFi, Oxford Nanopore Technologies and Hi-C reads (Figure 1). The assembly includes the W chromosome, which is notoriously difficult in birds because it is a refugium for repeats (Peona et al. 2021). We found that 28.34% of the snowy owl genome consists of repeats, which is one of the highest recorded for any bird and on par with that of the woodpeckers (Kapusta and Suh 2017; Manthey et al. 2018). The lion’s share of the repeats consists of a recent expansion of centromeric satellite repeats in Strigidae owls and is most likely the reason why the owl genomes are relatively large with C-values spanning from 1.4 to 2.0 (Animal genome size database, genomesize.com, accessed May 6th 2024). The snowy owl assembly was produced using two types of long reads (ONT and HiFi), which was crucial to resolve some of the complex repeat DNA regions, such as in the large regions of centromeric satellite repeats and the W chromosome (Figure 3). Currently neither technology results in complete representation of all DNA sequences. In chicken, ONT was better at resolving the euchromatic regions with high GC content on the smallest microchromosomes (‘dot chromosomes’) (Huang et al. 2023). Here we have identified some, but not all microchromosomes from unplaced scaffolds using both ONT and HiFi. Increasing ONT coverage in the future might amend this issue, as well as lead to placing more of the centromeric satellites in chromosomes.

Although the presence of centromeres is highly conserved in eukaryotes, the sequence is not, and it is now widely recognized that centromeres are determined epigenetically (Talbert and Henikoff 2022). The appearance of evolutionary new centromeres is therefore a widespread phenomenon, including in birds (Shang et al. 2010). A previous study has reported that the ancestor of the Strigidae family likely evolved new ENCs that replaced all previous centromere sequences, as indicated by the presence of lineage-specific centromere satellites on all chromosomes (Yamada et al. 2004). Our snowy owl genome assembly supports these previous findings and adds new insight into the evolution of the centromere position. We do not have the absolute timing of these repeat expansions. However, our finding that the putative centromere satellites (family-0 elements) are present in the snowy owl and not the barn owl genome supports that these satellite elements are lineage-specific to the Strigidae owls and are not present in Tytonidae. Family-0 could have been lost in the barn owl genome, however, it is reasonable to assume that this family-0 expansion happened after the split between Strigidae and Tytonidae ∼40 million years ago (Kuhl et al. 2021). Following the formation of ENCs in the snowy owl at least some of the centromeres have shifted position relative to chicken centromeric locations (Figure S8) and not all ENCs have evolved in fission breakpoints (Figs 4 and S6). To further elucidate the shifts in centromere position and role in chromosome evolution in aves requires centromere annotation for more avian genome assemblies.

A main result from our analyses of centromere-associated repeats is the notable sequence similarity between the centromere satellite DNA and ERV1 transposons. After the origin of novel centromere sequences, ENC matures over time by acquiring satellite DNA (Tolomeo et al. 2017), and in some cases, these satellites originate (i.e. are seeded) from a TE sequence. For instance, chicken ENCs are enriched for TEs compared with other centromeres (Shang et al. 2010), and in *Drosophila,* the functional centromere sequence that binds CENP-A is derived from a TE (Chang et al. 2019). Our results suggest that ENCs in the ancestor of the snowy owl were seeded by ERV1 transposons. The consensus sequence for the ERV1 family-0 element is 305 bp. There are typically 146 bp surrounding a nucleosome (van Holde 1989) and most centromeric repeat motifs are close to that size (Talbert and Henikoff 2022). However, in some cases, younger satellites have larger repeat motifs which can wrap two rounds around two nucleosomes forming a dimer (Henikoff et al. 2015). It therefore seems reasonable to believe that these ERV1s have taken on the function of centromeric satellite DNA. However, it is also possible that the ENCs in the snowy owl were later invaded by ERV1, erasing the traces of the initial satellite motifs. TE invasion is common in pericentromeric regions, such as in *Brassica rapa (Zhang et al. 2023)*, mice (Packiaraj and Thakur 2024), and in *Arabidopsis* species (Wlodzimierz et al. 2023). In the latter example, centromeres go through cycles of transposon invasion and satellite homogenization, creating complex higher-order repeats. Putatively, a similar process has shaped snowy owl centromeres given the two bursts of family-0 repeats (Figure S5B) and the presence of higher-order repeats in the putative centromeric regions (Figure 3B). Possibly the first burst reflects an initial period of ERV family-0 transposition, whereas the more recent copies of family-0 DNA originated from a subsequent expansion of satellite repeats seeded from the earlier ERV eruption. The region with highly identical family-0 elements represented as dark red triangles (Figure 3B) is most likely the functional centromere, with the more diverse repeats forming the pericentromeric region, but this would need to be confirmed by determining which motif in the genome that binds CENP-A and the rest of the kinetochore assembly. There are no other TEs in the centromere satellites, however the flanking regions contain multiple TEs (Figure 3B), including ERV1s. To distinguish between satellite seeding and invasion is difficult, however, given that these satellites are homologous between different species of Strigidae (Yamada et al. 2004), one seeding event seems more likely.

We have so far assumed selection is not the cause of the high similarity between the family-0 elements, but is rather due to a recent and neutral expansion of these satellite repeats. However, a recent study on the rapid evolution of new centromere satellites in the owl monkey posits the hypothesis that centromere replacement has been driven by adaptation to a nocturnal lifestyle (Nishihara et al. 2021). Satellites that previously had centromeric function have been co-opted to be part of heterochromatin blocks in the microlens of the eye. The possibility of satellite DNA being adaptive by having structural functions is a possible avenue of future research.

The observation of centromeric sequences that are highly similar within species, but divergent between species could also be explained by centromeric drive. Centromeres can behave as selfish genetic elements that compete to be one of the four meiotic products during female meiosis that becomes the final oocyte (Zwick et al. 1999). This centromeric drive selects for repeat expansions which can quickly be fixed in a population, but is counterbalanced by selection in the ‘host’ genome to prevent distortion of segregation during meiosis (Henikoff et al. 2001; Malik 2009). This evolutionary arms race can ultimately lead to diversification and speciation, which has been demonstrated in several model systems including chicken (Axelsson et al. 2010), the *Drosophila* genus (Ferree and Barbash 2009), and maize (reviewed by (Dawe 2022)). The presence of large centromeric satellite blocks that are highly similar in the snowy owl genome (Figure 2 and Figure S4) and phylogenetically within different species of Strigidae (Yamada et al. 2004) raises the question whether centromeric drive was involved in the diversification of this lineage. Similar patterns were reported in crows (Peona et al. 2023) and voles (Romanenko et al. 2018). Centromeric drive may therefore be quite prevalent in nature, but underreported as it requires pedigree data to track the fate of centromeres in female meiosis.

Rapid turnover of genomic architecture through ENC evolution, repeat expansions, and putative centromeric drive could lead to more frequent chromosome rearrangements and therefore cause breaks in collinearity with other species. However, we find that collinearity is highly conserved between the genomes of snowy owls and distant relatives such as chicken and zebra finch (Figure 4). This is concordant with a study showing that repositioning of centromeres in reptiles did not disrupt synteny with other vertebrates (Dobry et al. 2023). In contrast, the downy woodpecker genome is repeat-rich and highly rearranged compared with the other species in this study (Figure 3), including a fusion event between the Z chromosome and an autosome (chromosomal region homologous to chicken chromosome 1), which could be related to the recent high TE activity in this species (Manthey et al. 2018). Interchromosomal rearrangements between sex chromosomes and autosomes are rare events in birds, but have been documented before in other avian lineages (Sigeman et al. 2020; Huang et al. 2022). Clearly, TE bursts have had very different impacts on genome evolution in the snowy owl and downy woodpecker.

It is not unusual to have different centromeric sequences at different chromosomes in the same species, e.g. chickens have different centromeric satellite DNA at macro- and micro-chromosomes (Shang et al. 2010). Snowy owls seem to have similar, homologous satellites at all chromosomes (Yamada et al. 2004), which is confirmed in this study (Figure 3). Tytonidae, on the other hand, have a very different karyotype from other birds with no apparent distinction between macro- and micro-chromosomes (Rebholz et al. 1993; Yamada et al. 2004). The causes of this are currently unknown, and Tytonidae do not possess the specific group of ENC we observe in Strigidae nor have a particularly high repeat content (Figure 2).

The phylogenetic placement of Strigiformes within the Telluraves (core landbirds) has been debated over the years (Jarvis et al. 2014; Prum et al. 2015; Reddy et al. 2017; Braun et al. 2019), as well as the phylogenetic relationships within Strigiformes (Salter et al. 2020). Support for different topologies has depended on data type, method and/or number of taxa included in the analyses. A recent phylogenomic study of birds found that Strigiformes are phylogenetically closer to Accipitriformes than to Pittiformes + Coraciiformes (Stiller et al. 2024). This is concurrent with our results based on gene orthogroups (Figure 2 and Figure S6). Stiller et al. (2024) found that a sampling bias in favor of Passeriformes is the driver of not identifying Strigiformes and Accipitriformes as sister clades, highlighting the value of sequencing more species from these two groups. It is also plausible that the extensive repeat expansions in Pittiformes and Strigiformes could have resulted in long-branch attraction in previous phylogenetic studies. This could explain why topologies based purely on exons differ from topologies based partially or completely on noncoding regions (Reddy et al. 2017). The use of collinearity as a phylogenetic signal (Drillon et al. 2020) could be highly useful in future phylogenetic studies of birds. Given the highly conserved collinearity between Strigiformes and Accipitriformes (Figure 4), it seems parsimonious to suggest that they are indeed sister groups.

## Materials and Methods

### Genome Sample

We collected blood from a captive adult female snowy owl from the wildlife park Greifvogelstation & Wildfreigehege Hellenthal in Germany during a routine veterinary examination and put it on dry ice for preservation. Handling of the animal and blood collection were performed during routine hematological and blood chemical examinations and no additional handling, restrain or sampling was necessary for the purpose of this study. This individual was chosen for sequencing because obtaining a high-quality, snap-frozen tissue sample from a wild snowy owl is difficult in the wild. We further selected this individual because it is a female and thus has both sex chromosomes (W and Z, respectively). This snowy owl is captive-bred in Germany and believed to be of European ancestry.

### DNA sequencing

DNA isolation started from 10-20µl of ethanol-preserved blood per reaction. For ONT sequencing DNA extraction and library preparation was done using the Nanobind UL Library Prep Kit (Circulomics) with the combination of the Ultra-Long DNA Sequencing Kit (SQK-ULK001) (Oxford Nanopore) following the manufacture protocol. The library was loaded on an R9.4.1 flow cell and was sequenced using a Promethion24 device. To maximize the data output we performed a nuclease flush when the ratio of the sequencing pores dropped under 20%. Nuclease flush and library reloading were repeated twice. Guppy 5.1.13 was used for base calling with the High-accuracy base calling model.

For PacBio long-read sequencing, DNA extraction was performed according to the Circulomics protocol. First, ethanol was removed from the pelleted blood, and then washed twice in EtOH removal buffer (see Circulomics protocol). DNA was then isolated from the washed blood pellet using the Circulomics Nanobind CBB kit and protocol according to the manufacturer’s recommendations (Circulomics, now PacBio company). High molecular weight DNA was eluted from the Nanodisk with 150µl Tris-cl buffer, incubated overnight at room temperature before quality check of amount, purity, and integrity with UV-absorbance ratios (Nanodrop, Thermo Fisher), Qubit BR DNA quantification assay kit (Thermo Fisher) and Fragment Analyser using the DNA HS 50kb large fragment kit (Agilent Tech.) Before PacBio HiFi library preparation, DNA was purified for an additional time using PacBio Ampure Beads (1:1 ratio). Then 7.5µg of purified high molecular weight DNA was sheared to approx. 15-20 kb large fragments using the Megaruptor3 (Diagenode) in 200µl volume. We used 5µg of fragmented DNA with the PacBio library preparation kit 2.0 (PacBio). The final HiFi library was size-selected with a 10kb cut-off using a BluePippin instrument (Sage Biosciences) before sequencing on the PacBio Sequel II instrument at the Norwegian Sequencing Centre (NSC). We performed all QC steps with the same instrumentation as explained above.

HiC libraries were made using the Dovetail® Omni-C^®^ kit, following the Omni-C proximity Ligation Assay version 1.0 (Dovetail Inc), starting from 20-30µl of EtOH preserved blood. Final library quality was assayed using instrumentation described above as well as qPCR using the Kapa Library quantification kit for Illumina (Roche Inc.), before sequencing with other libraries on a quarter Illumina Novaseq S4 flowcell with 2*150 bp paired-end mode at the Norwegian Sequencing Centre.

### Genome Assembly and Curation

KMC v3.1.2rc1 was used to count k-mers of size 21 in the PacBio HiFi reads (Kokot et al. 2017). GenomeScope v2.0 was run on the k-mer histogram output from KMC to get estimates of genome size, heterozygosity, and repetitiveness (Ranallo-Benavidez et al. 2020). HiFiAdapterFilt v2.0.0 was applied on the HiFi reads to remove remnant PacBio adapter sequences (Sim et al. 2022). The filtered HiFi reads were assembled using hifiasm v0.16.1 with Hi-C integration (Cheng et al. 2022), resulting in a pair of haplotype-resolved assemblies, hap1, and hap2. Unique k-mers in each assembly were identified using meryl v1.3.0 (https://github.com/marbl/meryl), and used to create two sets of Hi-C reads, one without any k-mers occurring uniquely in hap1 and the other without k-mers occurring uniquely in hap2. The ONT reads were assembled with Flye 2.9-b1768 using default options (Kolmogorov et al. 2019). The Flye assembly was used to scaffold both haplotype-resolved assemblies from hifiasm using RagTag v2.1.0 (Alonge et al. 2021). K-mer-filtered Hi-C reads were aligned to each scaffolded assembly using BWA-MEM v0.7.17 with the −5SPM option (Li 2013). The alignments were sorted based on name using samtools 1.15.1, then samtools fixmate was applied, before default sorting and applying samtools markdup (Danecek et al. 2021). The resulting BAM file was used to further scaffold the two RagTag-scaffolded assemblies using YaHS v1.1a with default options (Zhou et al. 2023). The two resulting assemblies were then gap-filled using the Flye assembly with RagTag patch (Alonge et al. 2021).

FCS-Adaptor v0.2.2 (https://github.com/ncbi/fcs) was run on the assemblies, and any adaptor sequences found were masked using bedtools v2.30.0 maskfasta (Quinlan and Hall 2010). FCS-GX v0.2.2 (https://github.com/ncbi/fcs) was used to search for contamination. If a contaminant was found at the start or end of a sequence, the sequence was trimmed using a combination of samtools faidx, bedtools complement, and bedtools getfasta. If the contaminant was internal, it was masked using bedtools maskfasta. The mitochondrial genome was assembled using MitoHiFi v2.2, based on the contig assembly of pseudo-haplotype 1 (Uliano-Silva et al. 2023).

Merqury v1.3 was used to assess the completeness and quality of the genome assemblies by comparing them to the k-mer content of the Hi-C reads (Rhie et al. 2020). BUSCO v5.3.1 was used to assess the completeness of the genome assemblies by comparing them against the expected gene content in the Aves lineage (OrthoDB v10) (Manni, Berkeley, Seppey, and Zdobnov 2021). Gfastats v1.2.2 was used to output different assembly statistics of the assemblies (Formenti et al. 2022).

The assemblies were manually curated using the GRIT rapid curation suite(Howe et al. 2021) and the PretextView v.0.2.5 (https://github.com/wtsi-hpag/PretextView, last accessed October 25, 2022). During curation 119 and 89 corrections were made to pseudo-haplotypes 1 and 2, respectively. Through this, we identified 32 putative chromosomes for pseudo-haplotype 1, and 30 for pseudo-haplotype 2 via visual inspection. The Z and W chromosomes were curated by mapping against the chicken sex chromosomes using minimap v2.1 (Li 2021).

The Hi-C contact maps for both pseudo-haplotypes were visualized before and after curation using PretextSnapshot (https://github.com/wtsi-hpag/PretextSnapshot, last accessed October 25, 2022).

### Comparing pseudo-haplotypes 1 and 2

To compare the two pseudo-haplotype assemblies, we first aligned the homologous chromosomes using minimap2 v.2.1 (Li 2021) with the asm5 setting. We then used SyRI v1.6.3 (Goel et al. 2019) to identify structural variation between the two assemblies and visualized the results using plotsr v1.1.0 (Goel and Schneeberger 2022). For further comparisons, using the MUMmer toolkit v4.0.0rc1 (Marçais et al. 2018), we aligned the sequences using nucmer and then processed the alignment with dnadiff, which produced a list of indels, SNPs, and insertions all of which are reported in Table S1.

### Comparative Genomics Data

For comparative genomic analyses, we selected five species from the Telluraves clade (core landbirds, which includes Strigiformes) which have chromosome-level assemblies; downy woodpecker (*Dryobates pubescens*), Northern Carmine bee-eater (*Merops nubicus*), Northern goshawk (*Accipiter gentilis*), California condor (*Gymnogyps californianus*) and barn owl (*Tyto alba*). The latter species is not chromosome-level but was included as it is in the owl order (Strigiformes) and the only long-read assembly for owls available. We also included the genomes of chicken (*Gallus gallus*) and zebra finch (*Teaniopygia guttata*), as these are well-studied avian reference genomes. Versions of assemblies and an overview of genome characteristics are listed in Table S3.

### Genome Annotation

Gene annotations downloaded and used for the chicken, zebra finch, and California condor genomes are listed in Table S3. AGAT v1.0 agat_sp_keep_longest_isoform.pl and agat_sp_extract_sequences.pl were used on the chicken assembly and annotation to generate one protein (the longest isoform) per gene (Dainat et al. 2023). Miniprot v0.5 was used to align the proteins to the curated assemblies (Li 2023). UniProtKB/Swiss-Prot release 2022_03 (UniProt Consortium 2023) in addition to the Vertebrata part of OrthoDB v10 (Manni, Berkeley, Seppey, Simão, et al. 2021) were also aligned separately to the assemblies. RED v2018.09.10 was run via redmask (https://github.com/nextgenusfs/redmask) on the assemblies to mask repetitive areas (Girgis 2015). In addition RepeatModeler v2.0.3 (Flynn et al. 2020) was run on the hap1 genome assembly via the Dfam TE Tools container v1.6 (https://github.com/Dfam-consortium/TETools) to create a species-specific repeat library, and RepeatMasker 4.1.3-p1 (from the same container) was used to apply the library to the genome and discover repetitive elements (Smit et al. 2015). GALBA (https://github.com/Gaius-Augustus/GALBA, commit: f4aaeca) was run with the chicken proteins using the miniprot mode on the masked assemblies (Stanke et al. 2006; Buchfink et al. 2015; Hoff and Stanke 2019; Brůna et al. 2023). The funannotate-runEVM.py script from Funannotate v1.8.13 (Palmer and Stajich 2020) was used to run EvidenceModeler v1.1.1 (Haas et al. 2008) on the alignments of chicken proteins, UniProtKB/Swiss-Prot proteins, vertebrata proteins and the predicted genes from GALBA. The resulting predicted proteins were compared to the protein repeats that Funannotate distributes using DIAMOND v2.0.15 (Buchfink et al. 2015) blastp and the predicted genes were filtered based on this comparison using AGAT. The filtered proteins were compared to the UniProtKB/Swiss-Prot release 2022_03 using DIAMOND blastp to find gene names and InterProScan v5.47-82 (Jones et al. 2014) was used to discover functional domains. AGATs agat_sp_manage_functional_annotation.pl was used to attach the gene names and functional annotations to the predicted genes. For the Downy woodpecker, Northern Carmine bee-eater, Northern goshawk, and barn owl the same annotation pipeline was used as for the snowy owl, with the exception that we used the already soft-masked genome assemblies available at NCBI (Table S3).

### Synteny and orthology inference

To investigate chromosome-level synteny, including collinearity, between the snowy owl genome and chicken, zebra finch, California condor, Downy woodpecker, Northern Carmine bee-eater, Northern goshawk, and barn owl we used GENESPACE v0.9.3 (Lovell et al. 2022) with Orthofinder v2.5.4 (Emms and Kelly 2019) in default mode using gene annotations and protein sequences from each species. GENESPACE cannot handle too many scaffolds for comparing collinearity blocks, thus we included only chromosomes if chromosome information was available, or scaffolds >5Mb and putative chromosomes for the snowy owl (see information above). We ran Orthofinder separately on the entire dataset including unplaced scaffolds in order to assess genome-wide patterns of gene duplications/losses.

### Repeat landscape

We used the output from RepeatMasker (Smit et al. 2015) to plot the size distribution and percentage divergence from consensus using custom R scripts. We estimated the repeat density of different repeat classes in 200kb windows across the snowy owl genome using BEDTools v2.30.0 coverage (Quinlan and Hall 2010) and custom R script. We used StainedGlass (Vollger et al. 2022) to create identity heatmaps of tandem repeat structures in the putative centromere regions, with a window size of 2000 bp.

## Acknowledgements

The authors are grateful to the wildlife park Greifvogelstation & Wildfreigehege Hellenthal, Germany, for the supply of biological material for this study. We thank The International Snowy Owl Working Group (ISOWG) for advice and support. We thank the Vertebrate Genomes Project (VGP) for earlier access to the genomes generated, particularly the Carmine bee-eater genome (Thomas Gilbert) and the Downy Woodpecker genome (Matthew Fuxjager), and the Chair Erich Jarvis for his help. We are grateful to Luohao Xu for providing the centromere positions in the chicken genome assembly. All laboratory work was carried out at the Norwegian Sequencing Centre (NSC) and at the Centre for Integrative Genetics (CIGENE) laboratory. The computations were performed on resources provided by Sigma2 - the National Infrastructure for High-Performance Computing and Data Storage in Norway.

## Data Availability

The annotations, repeat masking and assemblies produced in this study are available from Zenodo at https://zenodo.org/records/12643816 (DOI: **10.5281/zenodo.12643816).**

## Author Contributions

The study was designed by HTB. Sampling was carried out by DF. DNA isolation, library preparation, and sequencing was carried out by AKT, MS, and MA. HTB, OKT, BGA, ELGE, SRS, KSJ, RN, SB, and AKK contributed to analyses and interpretation of results. Figures were made by HTB. Manuscript was written by HTB and OKT with input from all co-authors. All authors read and approved the final manuscript.

## References

1. Alonge M, Lebeigle L, Kirsche M, Aganezov S, Wang X, Lippman ZB, Schatz MC, Soyk S. 2021. Automated assembly scaffolding elevates a new tomato system for high-throughput genome editing. BioRxiv.

2. Altemose N, Logsdon GA, Bzikadze AV, Sidhwani P, Langley SA, Caldas GV, Hoyt SJ, Uralsky L, Ryabov FD, Shew CJ, et al. 2022. Complete genomic and epigenetic maps of human centromeres. Science 376:eabl4178.

3. Altschul SF, Gish W, Miller W, Myers EW, Lipman DJ. 1990. Basic local alignment search tool. J. Mol. Biol. 215:403–410.

4. Axelsson E, Albrechtsen A, van AP, Li L, Megens HJ, Vereijken ALJ, Crooijmans RPMA, Groenen MAM, Ellegren H, Willerslev E, et al. 2010. Segregation distortion in chicken and the evolutionary consequences of female meiotic drive in birds. Heredity 105:290–298.

5. Benham PM, Cicero C, Escalona M, Beraut E, Fairbairn C, Marimuthu MPA, Nguyen O, Sahasrabudhe R, King BL, Thomas WK, et al. 2023. Remarkably high repeat content in the genomes of sparrows: the importance of genome assembly completeness for transposable element discovery. BioRxiv.

6. Braun EL, Cracraft J, Houde P. 2019. Resolving the Avian Tree of Life from Top to Bottom: The Promise and Potential Boundaries of the Phylogenomic Era. In: Kraus RHS, editor. Avian Genomics in Ecology and Evolution: From the Lab into the Wild. Cham: Springer International Publishing. p. 151–210.

7. Brůna T, Li H, Guhlin J, Honsel D, Herbold S, Stanke M, Nenasheva N, Ebel M, Gabriel L, Hoff KJ. 2023. Galba: genome annotation with miniprot and AUGUSTUS. BMC Bioinformatics 24:327.

8. Buchfink B, Xie C, Huson DH. 2015. Fast and sensitive protein alignment using DIAMOND. Nat. Methods 12:59–60.

9. Chang C-H, Chavan A, Palladino J, Wei X, Martins NMC, Santinello B, Chen C-C, Erceg J, Beliveau BJ, Wu C-T, et al. 2019. Islands of retroelements are major components of Drosophila centromeres. PLoS Biol. 17:e3000241.

10. Cheng H, Jarvis ED, Fedrigo O, Koepfli K-P, Urban L, Gemmell NJ, Li H. 2022. Haplotype-resolved assembly of diploid genomes without parental data. Nat. Biotechnol. 40:1332–1335.

11. Dainat J, Hereñú D, Davis E, Crouch K, LucileSol, Agostinho N, Pascal-Git, Zollman Z, Tayyrov. 2023. NBISweden/AGAT: AGAT-v1.1.0. Zenodo.

12. Danecek P, Bonfield JK, Liddle J, Marshall J, Ohan V, Pollard MO, Whitwham A, Keane T, McCarthy SA, Davies RM, et al. 2021. Twelve years of SAMtools and BCFtools. Gigascience 10.

13. Dawe RK. 2022. The maize abnormal chromosome 10 meiotic drive haplotype: a review. Chromosome Res. 30:205–216.

14. Dobry J, Zhu Z, Zhou Q, Wapstra E, Deakin JE, Ezaz T. 2023. Fixed allele differences associated with the centromere reveal chromosome morphology and rearrangements in a reptile (Varanus acanthurus BOULENGER). Mol. Biol. Evol. 40.

15. Drillon G, Champeimont R, Oteri F, Fischer G, Carbone A. 2020. Phylogenetic reconstruction based on synteny block and gene adjacencies. Mol. Biol. Evol. 37:2747–2762.

16. Emms DM, Kelly S. 2019. OrthoFinder: phylogenetic orthology inference for comparative genomics. Genome Biol. 20:238.

17. Feng S, Stiller J, Deng Y, Armstrong J, Fang Q, Reeve AH, Xie D, Chen G, Guo C, Faircloth BC, et al. 2020. Dense sampling of bird diversity increases power of comparative genomics. Nature 587:252–257.

18. Ferree PM, Barbash DA. 2009. Species-specific heterochromatin prevents mitotic chromosome segregation to cause hybrid lethality in Drosophila. PLoS Biol. 7:e1000234.

19. Flynn JM, Hubley R, Goubert C, Rosen J, Clark AG, Feschotte C, Smit AF. 2020. RepeatModeler2 for automated genomic discovery of transposable element families. Proc Natl Acad Sci USA 117:9451–9457.

20. Formenti G, Abueg L, Brajuka A, Brajuka N, Gallardo-Alba C, Giani A, Fedrigo O, Jarvis ED. 2022. Gfastats: conversion, evaluation and manipulation of genome sequences using assembly graphs. Bioinformatics 38:4214–4216.

21. Girgis HZ. 2015. Red: an intelligent, rapid, accurate tool for detecting repeats de-novo on the genomic scale. BMC Bioinformatics 16:227.

22. Goel M, Schneeberger K. 2022. plotsr: visualizing structural similarities and rearrangements between multiple genomes. Bioinformatics 38:2922–2926.

23. Goel M, Sun H, Jiao W-B, Schneeberger K. 2019. SyRI: finding genomic rearrangements and local sequence differences from whole-genome assemblies. Genome Biol. 20:277.

24. Haas BJ, Salzberg SL, Zhu W, Pertea M, Allen JE, Orvis J, White O, Buell CR, Wortman JR. 2008. Automated eukaryotic gene structure annotation using EVidenceModeler and the Program to Assemble Spliced Alignments. Genome Biol. 9:R7.

25. Henikoff JG, Thakur J, Kasinathan S, Henikoff S. 2015. A unique chromatin complex occupies young α-satellite arrays of human centromeres. Sci. Adv. 1.

26. Henikoff S, Ahmad K, Malik HS. 2001. The centromere paradox: stable inheritance with rapidly evolving DNA. Science 293:1098–1102.

27. Hoff KJ, Stanke M. 2019. Predicting Genes in Single Genomes with AUGUSTUS. Curr. Protoc. Bioinformatics 65:e57.

28. van Holde KE. 1989. Chromatin. New York, NY: Springer New York

29. Howe K, Chow W, Collins J, Pelan S, Pointon D-L, Sims Y, Torrance J, Tracey A, Wood J. 2021. Significantly improving the quality of genome assemblies through curation. Gigascience 10.

30. Huang Z, De O Furo I, Liu J, Peona V, Gomes AJB, Cen W, Huang H, Zhang Y, Chen D, Xue T, et al. 2022. Recurrent chromosome reshuffling and the evolution of neo-sex chromosomes in parrots. Nat. Commun. 13:944.

31. Huang Z, Xu Z, Bai H, Huang Y, Kang N, Ding X, Liu J, Luo H, Yang C, Chen W, et al. 2023. Evolutionary analysis of a complete chicken genome. Proc Natl Acad Sci USA 120:e2216641120.

32. Jarvis ED, Mirarab S, Aberer AJ, Li B, Houde P, Li C, Ho SYW, Faircloth BC, Nabholz B, Howard JT, et al. 2014. Whole-genome analyses resolve early branches in the tree of life of modern birds. Science 346:1320–1331.

33. Jones P, Binns D, Chang H-Y, Fraser M, Li W, McAnulla C, McWilliam H, Maslen J, Mitchell A, Nuka G, et al. 2014. InterProScan 5: genome-scale protein function classification. Bioinformatics 30:1236–1240.

34. Kapusta A, Suh A. 2017. Evolution of bird genomes-a transposon’s-eye view. Ann. N. Y. Acad. Sci. 1389:164–185.

35. Kokot M, Dlugosz M, Deorowicz S. 2017. KMC 3: counting and manipulating k-mer statistics. Bioinformatics 33:2759–2761.

36. Kolmogorov M, Yuan J, Lin Y, Pevzner PA. 2019. Assembly of long, error-prone reads using repeat graphs. Nat. Biotechnol. 37:540–546.

37. Kuhl H, Frankl-Vilches C, Bakker A, Mayr G, Nikolaus G, Boerno ST, Klages S, Timmermann B, Gahr M. 2021. An Unbiased Molecular Approach Using 3’-UTRs Resolves the Avian Family-Level Tree of Life. Mol. Biol. Evol. 38:108–127.

38. Li H. 2013. Aligning sequence reads, clone sequences and assembly contigs with BWA-MEM. arXiv.

39. Li H. 2021. New strategies to improve minimap2 alignment accuracy. Bioinformatics 37:4572–4574.

40. Li H. 2023. Protein-to-genome alignment with miniprot. Bioinformatics 39.

41. Lovell JT, Sreedasyam A, Schranz ME, Wilson M, Carlson JW, Harkess A, Emms D, Goodstein DM, Schmutz J. 2022. GENESPACE tracks regions of interest and gene copy number variation across multiple genomes. eLife 11.

42. Malik HS. 2009. The centromere-drive hypothesis: a simple basis for centromere complexity. Prog. Mol. Subcell. Biol. 48:33–52.

43. Manni M, Berkeley MR, Seppey M, Simão FA, Zdobnov EM. 2021. BUSCO Update: Novel and Streamlined Workflows along with Broader and Deeper Phylogenetic Coverage for Scoring of Eukaryotic, Prokaryotic, and Viral Genomes. Mol. Biol. Evol. 38:4647–4654.

44. Manni M, Berkeley MR, Seppey M, Zdobnov EM. 2021. BUSCO: assessing genomic data quality and beyond. Curr. Protoc. 1:e323.

45. Manthey JD, Moyle RG, Boissinot S. 2018. Multiple and independent phases of transposable element amplification in the genomes of piciformes (woodpeckers and allies). Genome Biol. Evol. 10:1445–1456.

46. Marçais G, Delcher AL, Phillippy AM, Coston R, Salzberg SL, Zimin A. 2018. MUMmer4: A fast and versatile genome alignment system. PLoS Comput. Biol. 14:e1005944.

47. Nishihara H, Stanyon R, Tanabe H, Koga A. 2021. Replacement of owl monkey centromere satellite by a newly evolved variant was a recent and rapid process. Genes Cells 26:979–986.

48. Ola M, O’Brien CE, Coughlan AY, Ma Q, Donovan PD, Wolfe KH, Butler G. 2020. Polymorphic centromere locations in the pathogenic yeast Candida parapsilosis. Genome Res. 30:684–696.

49. Packiaraj J, Thakur J. 2024. DNA satellite and chromatin organization at mouse centromeres and pericentromeres. Genome Biol. 25:52.

50. Palmer JM, Stajich J. 2020. Funannotate v1.8.1: Eukaryotic genome annotation. Zenodo.

51. Peona V, Kutschera VE, Blom MPK, Irestedt M, Suh A. 2023. Satellite DNA evolution in Corvoidea inferred from short and long reads. Mol. Ecol. 32:1288–1305.

52. Peona V, Palacios-Gimenez OM, Blommaert J, Liu J, Haryoko T, Jønsson KA, Irestedt M, Zhou Q, Jern P, Suh A. 2021. The avian W chromosome is a refugium for endogenous retroviruses with likely effects on female-biased mutational load and genetic incompatibilities. Philos. Trans. R. Soc. Lond. B Biol. Sci. 376:20200186.

53. Prum RO, Berv JS, Dornburg A, Field DJ, Townsend JP, Lemmon EM, Lemmon AR. 2015. A comprehensive phylogeny of birds (Aves) using targeted next-generation DNA sequencing. Nature 526:569–573.

54. Quinlan AR, Hall IM. 2010. BEDTools: a flexible suite of utilities for comparing genomic features. Bioinformatics 26:841–842.

55. Ranallo-Benavidez TR, Jaron KS, Schatz MC. 2020. GenomeScope 2.0 and Smudgeplot for reference-free profiling of polyploid genomes. Nat. Commun. 11:1432.

56. Rebholz WER, Boer LEMD, Sasaki M, Belterman RHR, Nishida-Umehara C. 1993. The chromosomal phylogeny of owls (strigiformes) and new karyotypes of seven species. Cytologia (Tokyo) 58:403–416.

57. Reddy S, Kimball RT, Pandey A, Hosner PA, Braun MJ, Hackett SJ, Han K-L, Harshman J, Huddleston CJ, Kingston S, et al. 2017. Why Do Phylogenomic Data Sets Yield Conflicting Trees? Data Type Influences the Avian Tree of Life more than Taxon Sampling. Syst. Biol. 66:857–879.

58. Rhie A, Walenz BP, Koren S, Phillippy AM. 2020. Merqury: reference-free quality, completeness, and phasing assessment for genome assemblies. Genome Biol. 21:245.

59. Romanenko SA, Serdyukova NA, Perelman PL, Trifonov VA, Golenishchev FN, Bulatova NS, Stanyon R, Graphodatsky AS. 2018. Multiple intrasyntenic rearrangements and rapid speciation in voles. Sci. Rep. 8:14980.

60. Salter JF, Oliveros CH, Hosner PA, Manthey JD, Robbins MB, Moyle RG, Brumfield RT, Faircloth BC. 2020. Extensive paraphyly in the typical owl family (Strigidae). Auk 137.

61. Sasaki M, Nishida-Umehara C, Tsuchiya K. 1994. A Comparative Study of G-banded Karyotypes in Eight Species of Owls. Cytologia (Tokyo) 59:183–185.

62. Shang W-H, Hori T, Toyoda A, Kato J, Popendorf K, Sakakibara Y, Fujiyama A, Fukagawa T. 2010. Chickens possess centromeres with both extended tandem repeats and short non-tandem-repetitive sequences. Genome Res. 20:1219–1228.

63. Sigeman H, Ponnikas S, Hansson B. 2020. Whole-genome analysis across 10 songbird families within Sylvioidea reveals a novel autosome-sex chromosome fusion. Biol. Lett. 16:20200082.

64. Sim SB, Corpuz RL, Simmonds TJ, Geib SM. 2022. HiFiAdapterFilt, a memory efficient read processing pipeline, prevents occurrence of adapter sequence in PacBio HiFi reads and their negative impacts on genome assembly. BMC Genomics 23:157.

65. Smalec BM, Heider TN, Flynn BL, O’Neill RJ. 2019. A centromere satellite concomitant with extensive karyotypic diversity across the Peromyscus genus defies predictions of molecular drive. Chromosome Res. 27:237–252.

66. Smit AFA, Hubley R, Green P. 2015. RepeatMasker Open-4.0. RepeatMasker Open-4.0 [Internet]. Available from: http://www.repeatmasker.org/

67. Stanke M, Schöffmann O, Morgenstern B, Waack S. 2006. Gene prediction in eukaryotes with a generalized hidden Markov model that uses hints from external sources. BMC Bioinformatics 7:62.

68. Stiller J, Feng S, Chowdhury A-A, Rivas-González I, Duchêne DA, Fang Q, Deng Y, Kozlov A, Stamatakis A, Claramunt S, et al. 2024. Complexity of avian evolution revealed by family-level genomes. Nature 629:851–860.

69. Talbert PB, Henikoff S. 2022. The genetics and epigenetics of satellite centromeres. Genome Res. 32:608–615.

70. Tolomeo D, Capozzi O, Stanyon RR, Archidiacono N, D’Addabbo P, Catacchio CR, Purgato S, Perini G, Schempp W, Huddleston J, et al. 2017. Epigenetic origin of evolutionary novel centromeres. Sci. Rep. 7:41980.

71. Uliano-Silva M, Ferreira JGRN, Krasheninnikova K, Darwin Tree of Life Consortium, Formenti G, Abueg L, Torrance J, Myers EW, Durbin R, Blaxter M, et al. 2023. MitoHiFi: a python pipeline for mitochondrial genome assembly from PacBio high fidelity reads. BMC Bioinformatics 24:288.

72. UniProt Consortium. 2023. Uniprot: the universal protein knowledgebase in 2023. Nucleic Acids Res. 51:D523–D531.

73. Vollger MR, Kerpedjiev P, Phillippy AM, Eichler EE. 2022. StainedGlass: interactive visualization of massive tandem repeat structures with identity heatmaps. Bioinformatics 38:2049–2051.

74. Waters PD, Patel HR, Ruiz-Herrera A, Álvarez-González L, Lister NC, Simakov O, Ezaz T, Kaur P, Frere C, Grützner F, et al. 2021. Microchromosomes are building blocks of bird, reptile, and mammal chromosomes. Proc Natl Acad Sci USA 118.

75. Wells JN, Feschotte C. 2020. A field guide to eukaryotic transposable elements. Annu. Rev. Genet. 54:539–561.

76. Wlodzimierz P, Rabanal FA, Burns R, Naish M, Primetis E, Scott A, Mandáková T, Gorringe N, Tock AJ, Holland D, et al. 2023. Cycles of satellite and transposon evolution in Arabidopsis centromeres. Nature.

77. Xu L, Auer G, Peona V, Suh A, Deng Y, Feng S, Zhang G, Blom MPK, Christidis L, Prost S, et al. 2019. Dynamic evolutionary history and gene content of sex chromosomes across diverse songbirds. Nat. Ecol. Evol. 3:834–844.

78. Xu L, Zhou Q. 2020. The Female-Specific W Chromosomes of Birds Have Conserved Gene Contents but Are Not Feminized. Genes 11.

79. Yamada K, Nishida-Umehara C, Matsuda Y. 2004. A new family of satellite DNA sequences as a major component of centromeric heterochromatin in owls (Strigiformes). Chromosoma 112:277–287.

80. Zhang G, Li C, Li Q, Li B, Larkin DM, Lee C, Storz JF, Antunes A, Greenwold MJ, Meredith RW, et al. 2014. Comparative genomics reveals insights into avian genome evolution and adaptation. Science 346:1311–1320.

81. Zhang L, Liang J, Chen H, Zhang Z, Wu J, Wang X. 2023. A near-complete genome assembly of Brassica rapa provides new insights into the evolution of centromeres. Plant Biotechnol. J. 21:1022–1032.

82. Zhou C, McCarthy SA, Durbin R. 2023. YaHS: yet another Hi-C scaffolding tool. Bioinformatics 39.

83. Zwick ME, Salstrom JL, Langley CH. 1999. Genetic variation in rates of nondisjunction: association of two naturally occurring polymorphisms in the chromokinesin nod with increased rates of nondisjunction in Drosophila melanogaster. Genetics 152:1605–1614.

