## Supplementary information for "Evolutionary new centromeres in the snowy owl genome putatively seeded from a transposable element"

### Supporting Information

#### Supporting figures

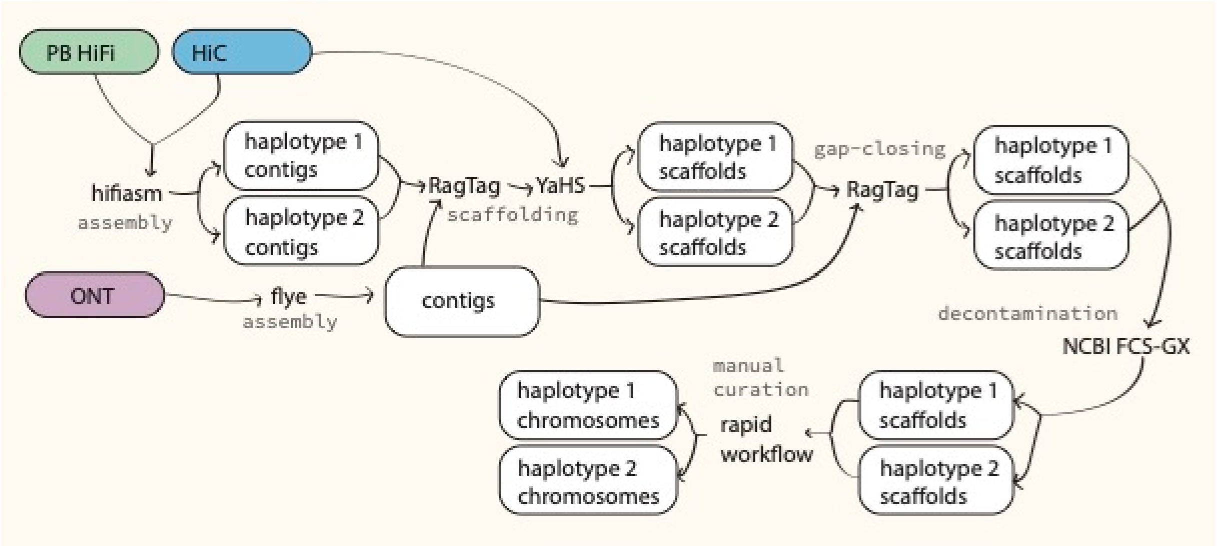


Figure S1: Assembly pipeline for the snowy owl genome version bBubSca1 presented in this study.

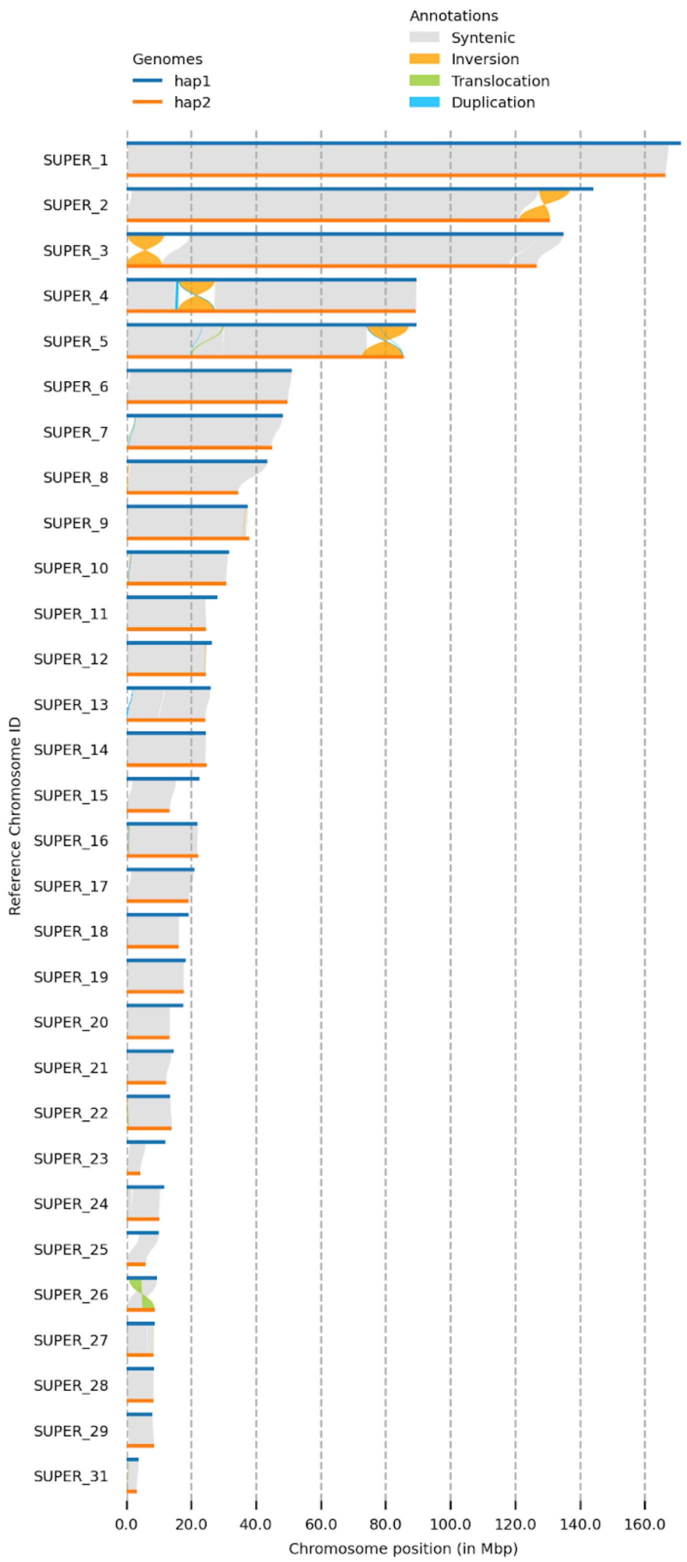


Figure S2: Synteny and Rearrangement Identifier (SyRi) plot comparing 30 chromosomes from pseudo-haplotype 1 and pseudo-haplotype 2 of the snowy owl genome assembly. The gray blocks show areas that are aligned, while white areas represent unaligned regions. Genomic rearrangements (i.e. inversions, duplications, and translocations) between the two pseudo-haplotypes are indicated according to legend.

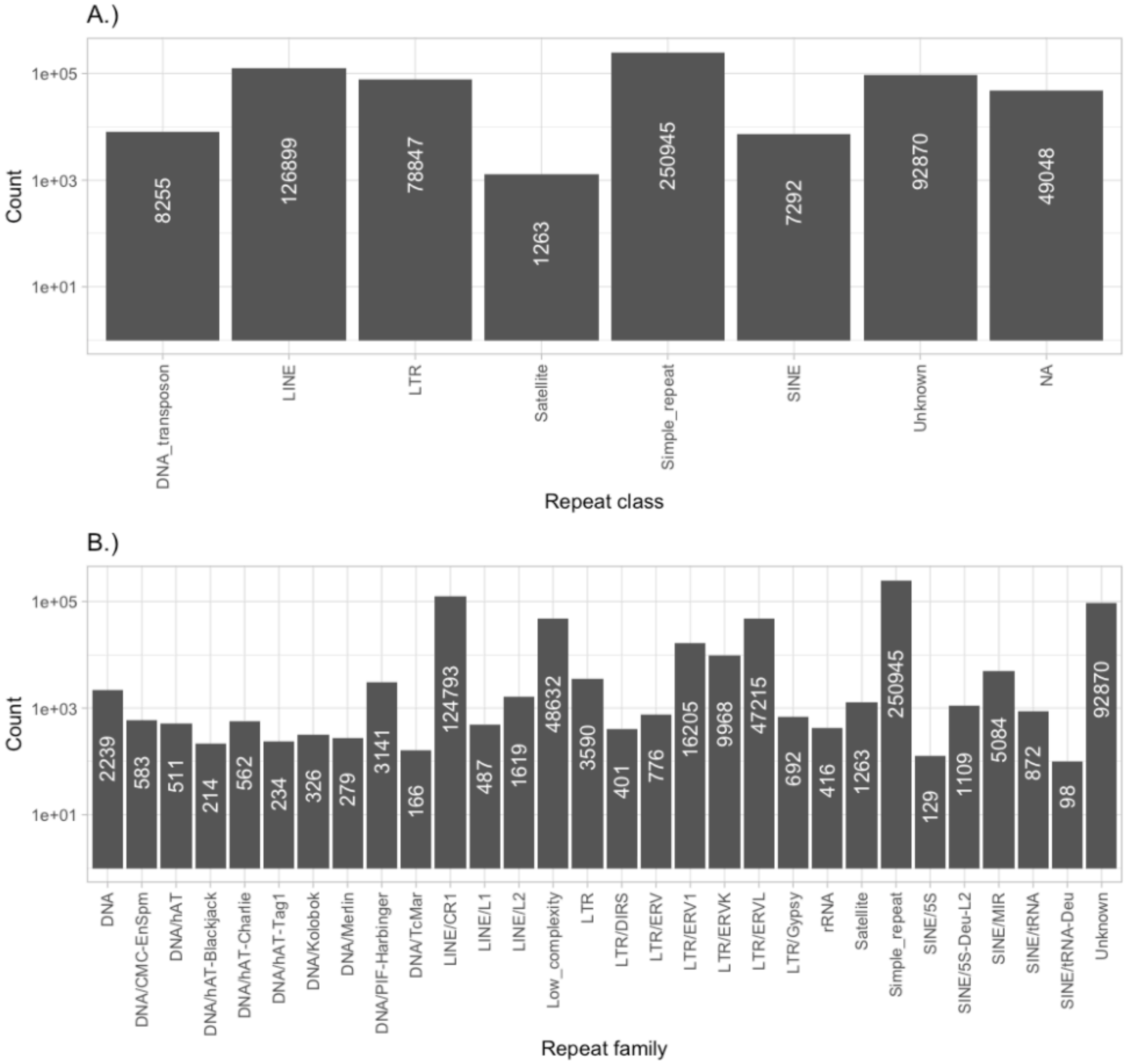


Figure S3: Counts of different types of repeats at a log10 scale in the snowy owl hap1 assembly at A.) class level and B.) family level.

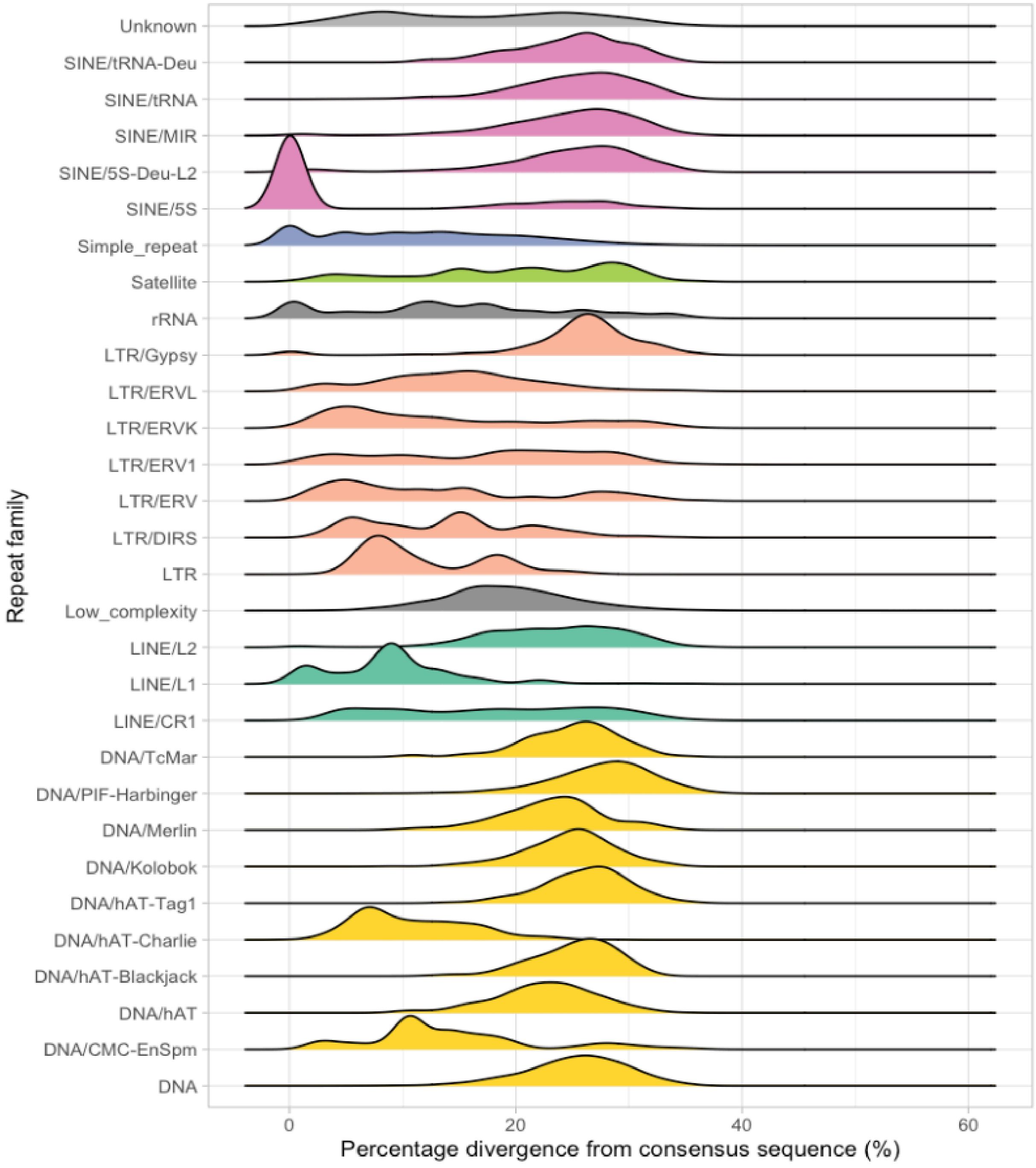


Figure S4: Percentage divergence from the consensus of repeats in the snowy owl genome for each repeat family.

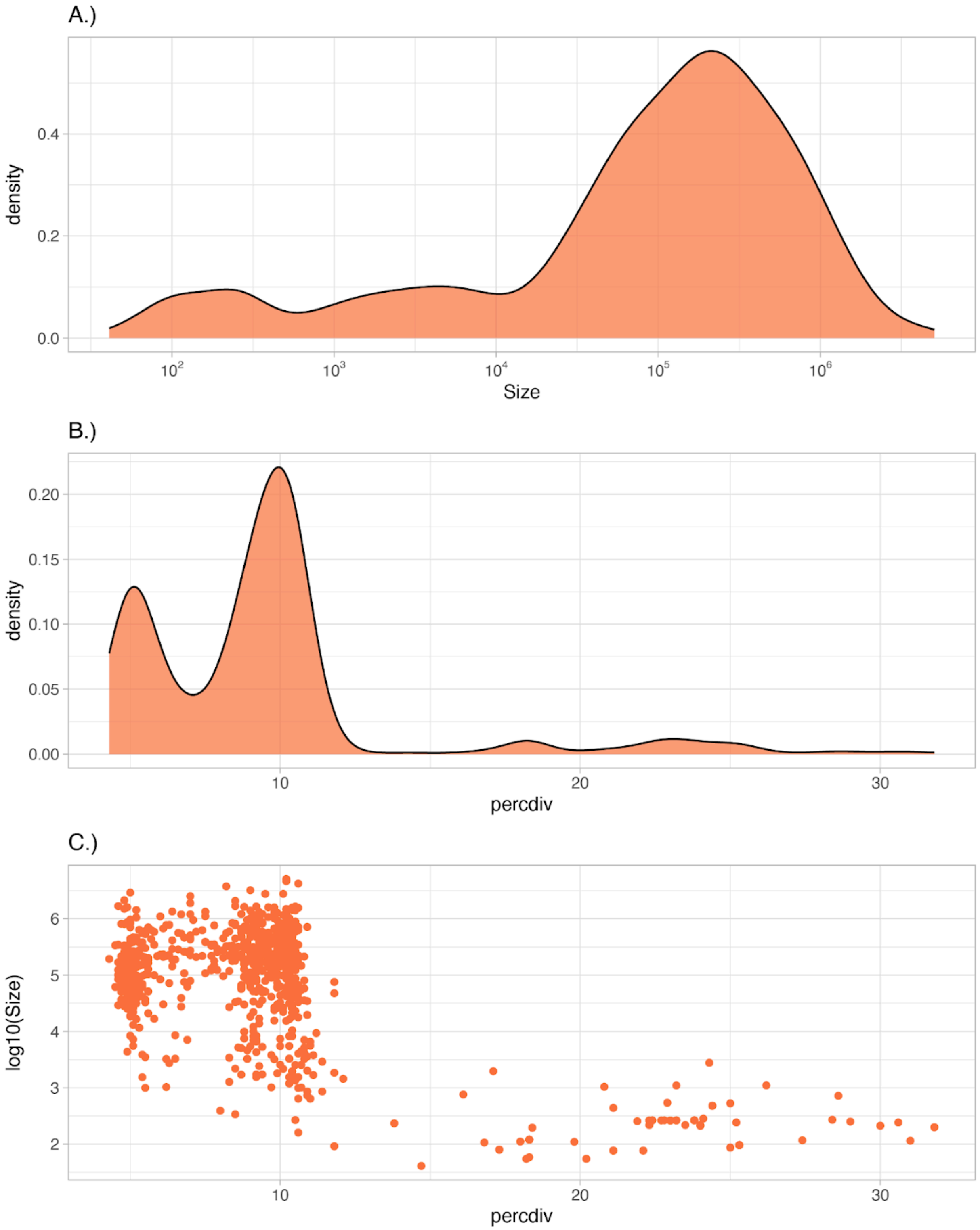


Figure S5: Statistics for the repeat type family family-0 (number of annotated family-0 elements = 16 205) in the snowy owl hap1: A.) Size distribution, B.) percentage divergence from consensus and C.) the relationship between size (log10 scale) and percentage divergence from consensus.

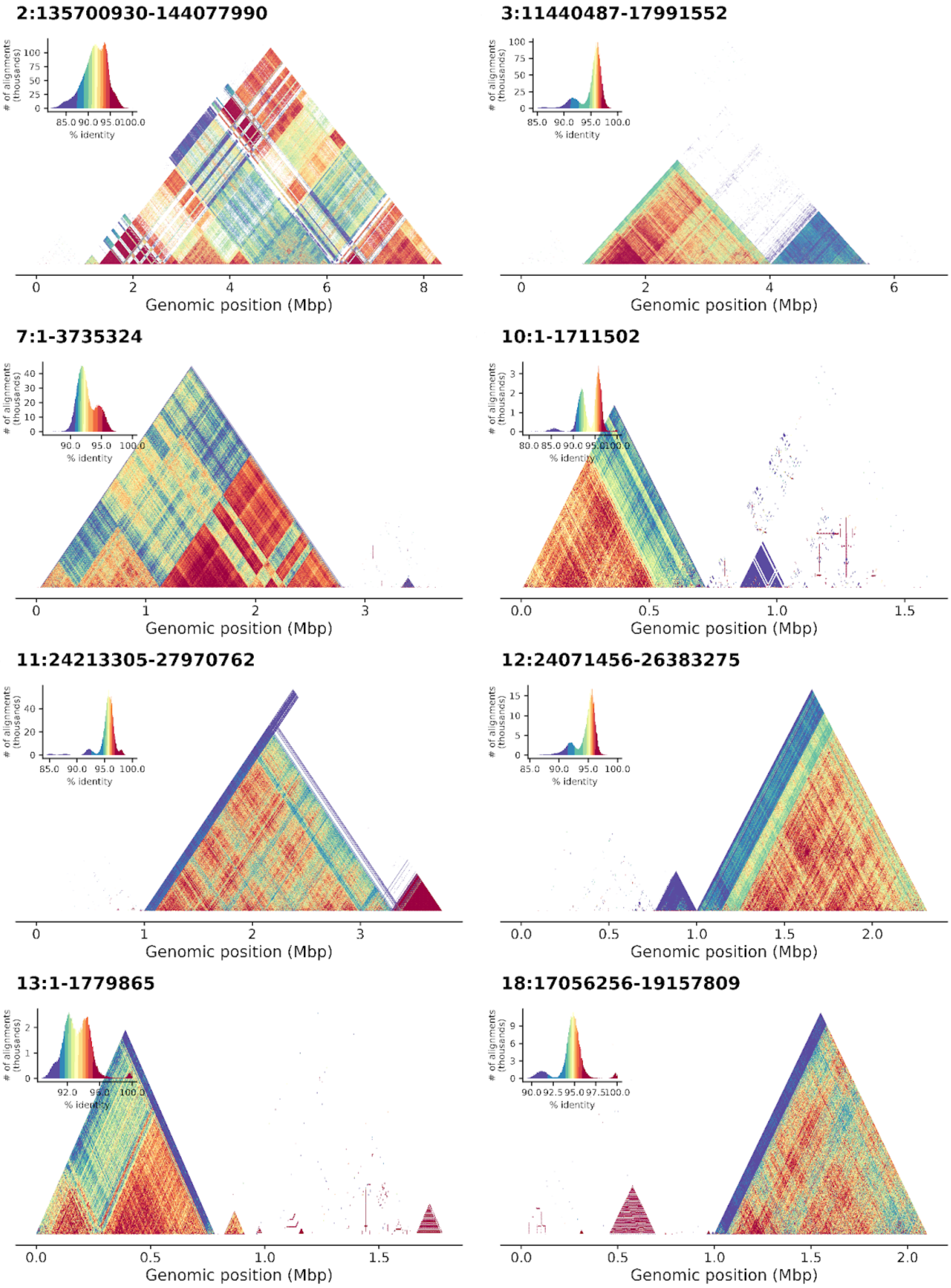

2:135700930-144077990
3:11440487-17991552
7:1-3735324
10:1-1711502
11:24213305-27970762
12:24071456-26383275
13:1-1779865
18:17056256-19157809
### of alignments (thousands)
% identity
Genomic position (Mbp)

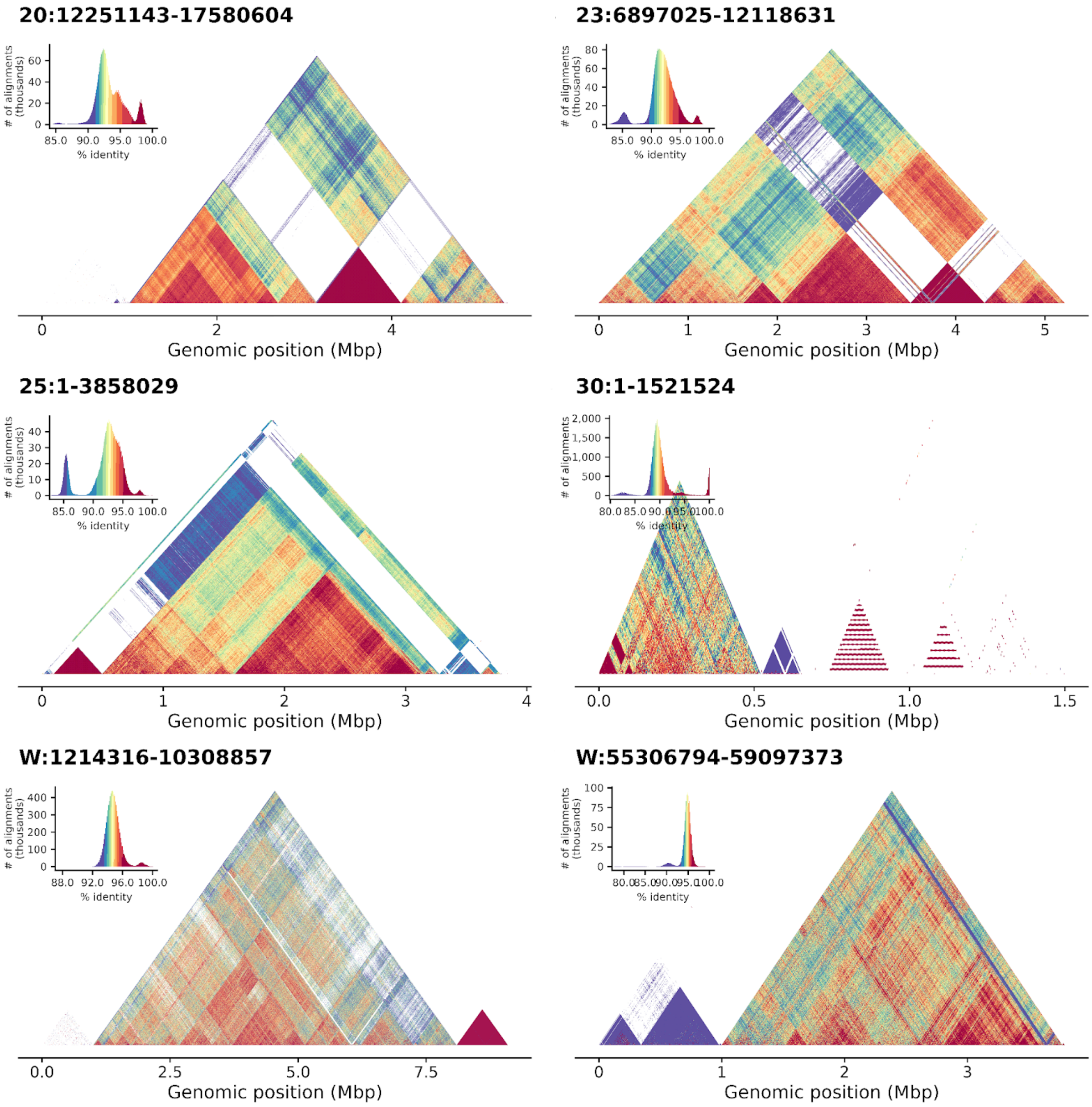


Figure S6: Examples of higher-order repeat structure of the putative centromere/pericentromere regions for all chromosomes with putative centromeres in the assembly. Each region is delimited by the position of family-0 elements in addition to 1 Mb of flanking sequence on either side (unless this exceeds the start or end of the chromosome, then the start or end is the limit). The heatmaps display an all-by-all percentage identity colored according to legend, calculated for 2 kb windows using StainedGlass.

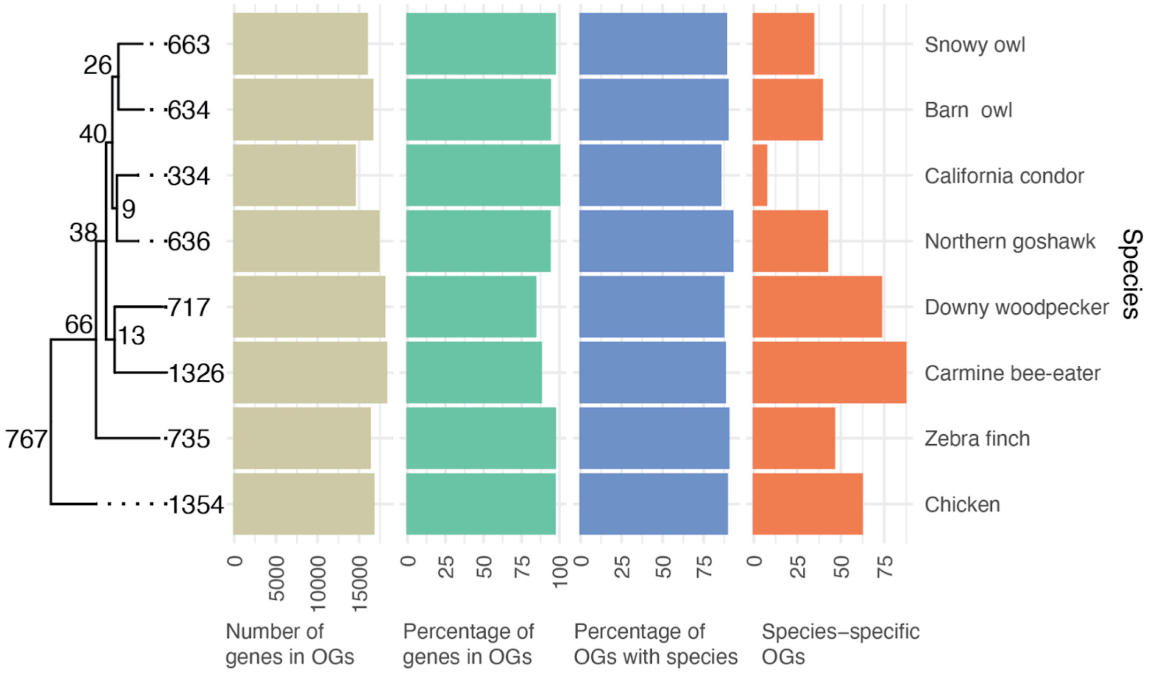


Figure S7: Orthofinder results for the eight bird species in this study. The number of gene duplications is denoted at each tip and node. Barplots show the number of genes placed in orthogroups (OGs), the percentage of genes in orthogroups, the percentage of orthogroups containing that species, and how many species-specific orthogroups there are.

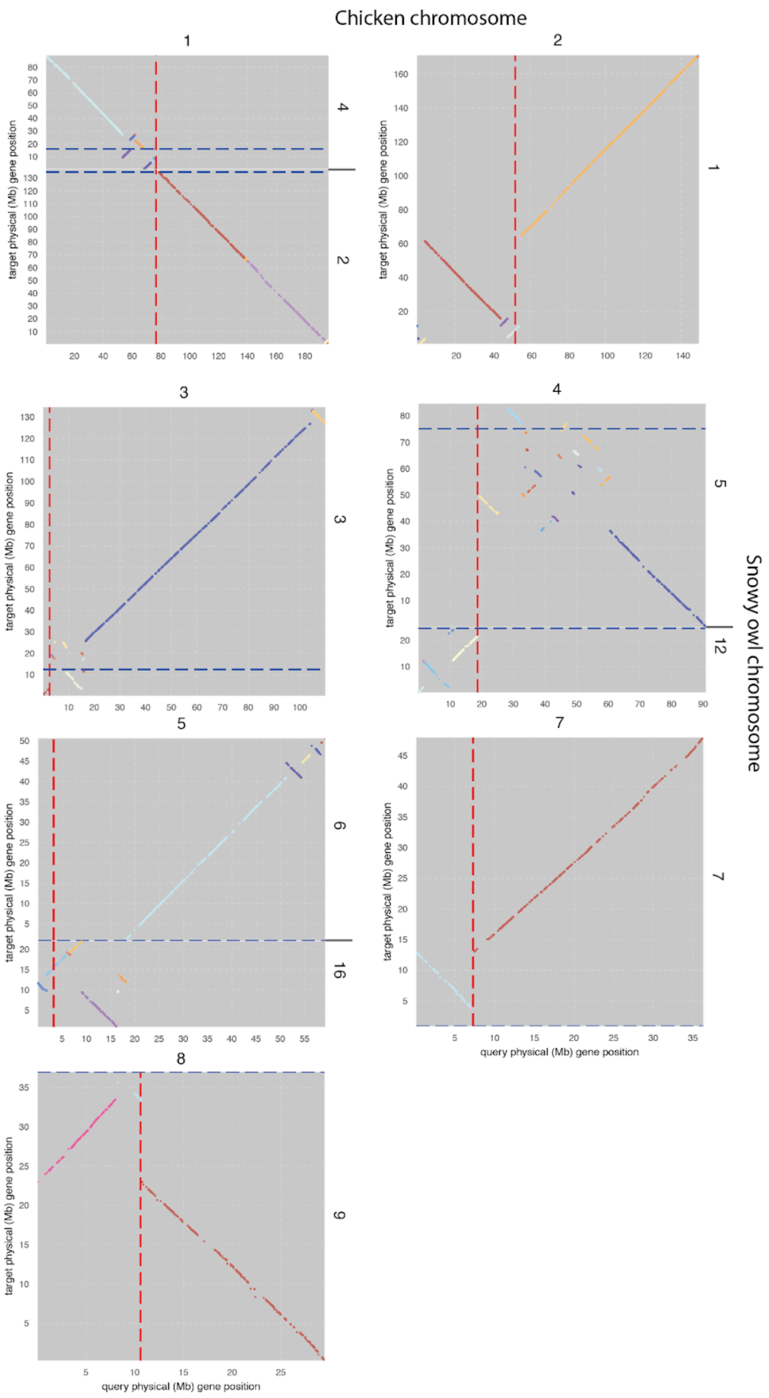


Figure S8: Dot plots showing collinearity between metacentric/submetacentric/subtelocentric chicken chromosomes and snowy owl chromosomes. Red dotted lines are the centromere position in chicken (Huang et al. 2023), blue dotted line is the start point of family-0 repeat chunk indicating putative centromere position in snowy owl, cross-referenced with cytogenetic results from (Yamada et al. 2004). Centromere positions for chromosomes 1 and 16 in the snowy owl are not known, however, they are most likely acrocentric (Yamada et al. 2004). Different colors indicate different collinearity blocks from GENESPACE.

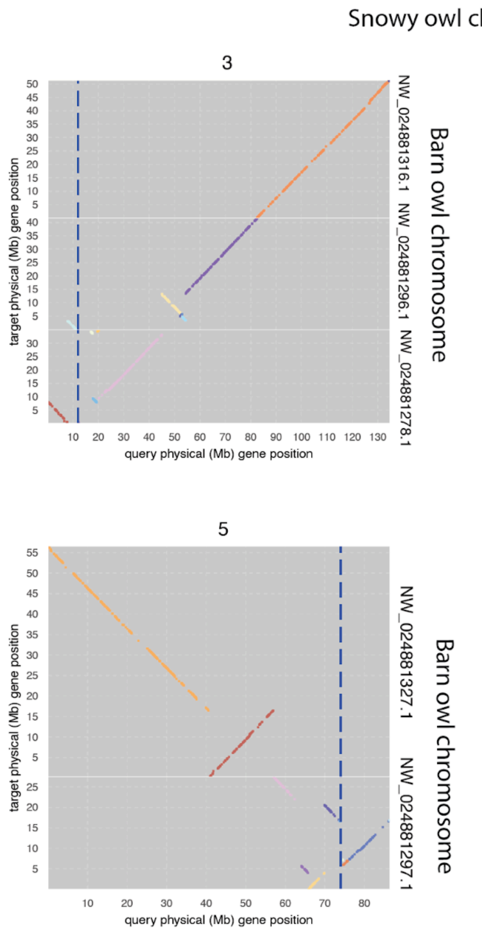


Figure S9: Dot plots between metacentric/submetacentric/subtelocentric snowy owl chromosomes and syntenic breakpoints in barn owl. Blue dotted lines are putative centromere positions in the snowy owl based on the position of family-0 repeats cross-referenced with cytogenetic results from (Yamada et al. 2004). Different colors indicate different synteny blocks from GENESPACE.

### Supporting tables

Table S1:Genome assembly statistics for *Bubo scandiacus*, bBubSca1.

| | | |
|---|---|---|
| **Project accession data** | | |
| Species | *Bubo scandiacus* | |
| Specimen | bBubSca1 | |
| Isolate information | Female, blood | |
| **Raw data accessions** | | |
| PacBio HiFi reads | 3 PACBIO_SMRT (Sequel II) runs: 3.6 M reads, 55.5 Gbp | |
| Oxford Nanopore reads | Sequenced on one FLO-PRO002 promethion flow cell, filtered 969 k reads, 19.4 Gbp | |
| Hi-C Illumina reads | 1 ILLUMINA (Illumina NovaSeq S4) run: 436 M pairs of reads, 131.7 Gbp | |
| **Genome assembly metrics** | | |
| HiFi read coverage | 39x | |
| Assembly identifier | bBubSca1.1.hap1 | bBubSca1.1.hap2 |
| Span (Mbp) | 1604 | 1233 |
| Number of contigs | 1430 | 842 |
| Contig N50 length (Mbp) | 16.5 | 21.4 |
| Number of scaffolds | 1105 | 639 |
| Scaffold N50 length (Mbp) | 51.0 | 49.8 |
| Longest scaffold (Mbp) | 171.1 | 166.5 |
| Consensus quality (QV) compared to Hi-C (compared to HiFi) | 39.6566 (47.7734) | 39.6335 (49.6791) |
| Both assemblies | 39.6465 (48.5017) | |
| *k*-mer completeness (percentage; compared to HiFi) | 97.3694 (97.8052) | 87.5326 (87.9751) |

| | | | |
|---|---|---|---|
| Both assemblies | | 98.9431 (99.4783) | |
| BUSCO* | | C:97.1%[S:96.1%,D:1.0%],F:0.5%, M:2.4%,n:8338 | C:89.9%[S:89.4%,D:0.5%],F:0.6%, M:9.5%,n:8338 |
| Percentage of assembly mapped to chromosomes | | 83.16 | 87.61 |
| Flagger** | | H: 80.23%, D: 16.82%, E:1.21%, C:1.75% | H: 86.06%, D: 12.13%, E:1.46%, C:0.34% |
| mummer | aligned bases | 1,102,853,948 (94.5016%) | 1,077,569,433 (99.7106%) |
| | Insertions (sum in bp) | 5422 (106,446,887) | 2483 (12,796,059) |
| | SNPs | 891827 | 891827 |
| | Indels | 592002 | 592002 |
| Sex chromosomes | | ZW | |
| Organelles | | (not identified) | (not identified) |
| **Genome annotation** | | | |
| Number of protein-coding genes | | 16955 | 15947 |
| BUSCO* | | C:96.5%[S:95.6%,D:0.9%],F:0.4%, M:3.1%,n:8338 | C:89.9%[S:89.4%,D:0.5%],F:0.6%, M:9.5%,n:8338 |

* BUSCO scores based on the aves BUSCO set using v5.4.7. C = complete [S = single copy, D = duplicated], F = fragmented, M = missing, n = number of orthologues in comparison.

**Flagger scores H = haploid, D = duplicated, E = error, C = collapsed

Table S2: Chromosomal pseudomolecules in the genome assembly of *Bubo scandiacus*, bBubSca1, hap1, and hap2.

| | Hap1 | | | | Hap2 | | | |
|---|---|---|---|---|---|---|---|---|
| Chromosome | Size (Mb) | GC% | Hap% | Dup% | Size (Mb) | GC% | Hap% | Dup% |
| 1 | 171.10 | 40.16 | 93.21 | 5.49 | 166.47 | 39.90 | 93.96 | 5.31 |
| 2 | 144.08 | 40.94 | 87.84 | 9.28 | 130.74 | 40.30 | 93.16 | 5.99 |
| 3 | 134.86 | 41.04 | 93.15 | 4.95 | 126.67 | 40.44 | 93.86 | 5.08 |
| 4 | 89.47 | 41.12 | 93.59 | 5.66 | 89.39 | 41.13 | 92.96 | 6.06 |
| 5 | 89.42 | 39.93 | 92.14 | 7.13 | 85.63 | 39.97 | 91.91 | 7.36 |
| 6 | 50.97 | 41.44 | 90.43 | 5.01 | 49.84 | 41.27 | 89.14 | 4.73 |
| 7 | 48.34 | 41.99 | 89.49 | 8.78 | 44.92 | 41.37 | 93.45 | 6.03 |
| 8 | 43.56 | 41.80 | 83.03 | 3.85 | 34.43 | 41.56 | 94.05 | 5.02 |
| 9 | 37.48 | 41.85 | 93.50 | 6.07 | 37.90 | 41.97 | 92.58 | 7.10 |
| 10 | 31.66 | 42.77 | 91.63 | 7.85 | 30.74 | 42.48 | 93.73 | 5.70 |
| 11 | 27.97 | 43.79 | 91.83 | 5.43 | 24.61 | 42.92 | 93.62 | 5.90 |
| 12 | 26.38 | 43.58 | 90.39 | 8.01 | 24.56 | 42.97 | 92.59 | 5.93 |
| 13 | 26.08 | 43.37 | 85.81 | 5.80 | 24.36 | 42.94 | 89.70 | 4.73 |
| 14 | 24.57 | 42.28 | 92.00 | 6.34 | 24.75 | 42.29 | 90.18 | 7.11 |
| 15 | 22.48 | 44.37 | 75.58 | 2.88 | 13.40 | 42.84 | 94.20 | 5.20 |
| 16 | 22.00 | 42.27 | 93.43 | 6.02 | 22.09 | 42.37 | 93.13 | 6.42 |
| 17 | 20.98 | 44.66 | 92.92 | 6.73 | 19.30 | 44.27 | 94.06 | 5.55 |
| 18 | 19.16 | 45.79 | 86.07 | 5.54 | 16.17 | 44.81 | 81.02 | 7.90 |
| 19 | 18.35 | 45.71 | 89.94 | 6.22 | 17.70 | 45.50 | 91.14 | 6.18 |
| 20 | 17.58 | 48.40 | 90.04 | 9.50 | 13.40 | 47.69 | 89.84 | 9.69 |
| 21 | 14.65 | 46.86 | 91.31 | 7.35 | 12.20 | 46.04 | 90.18 | 8.97 |

| | | | | | | | | |
|---|---|---|---|---|---|---|---|---|
| 22 | 13.55 | 46.61 | 93.24 | 6.23 | 13.96 | 46.75 | 90.84 | 8.60 |
| 23 | 12.12 | 51.93 | 87.21 | 9.79 | 4.29 | 50.48 | 81.92 | 16.00 |
| 24 | 11.64 | 47.42 | 79.44 | 19.00 | 10.17 | 47.07 | 84.41 | 14.73 |
| 25 | 10.03 | 49.66 | 84.96 | 14.25 | 6.04 | 48.84 | 80.06 | 19.27 |
| 26 | 9.55 | 50.26 | 81.80 | 17.60 | 8.68 | 50.37 | 86.65 | 12.88 |
| 27 | 8.69 | 51.71 | 72.21 | 26.27 | 8.27 | 51.62 | 77.03 | 21.14 |
| 28 | 8.54 | 49.03 | 82.49 | 17.02 | 8.38 | 49.00 | 83.85 | 15.68 |
| 29 | 8.03 | 51.65 | 83.24 | 12.90 | 8.47 | 51.38 | 70.87 | 14.78 |
| 30 | 4.65 | 56.40 | 69.42 | 19.57 | - | - | - | - |
| 31 | 3.74 | 57.02 | 47.03 | 44.70 | 3.16 | 59.13 | 43.31 | 40.01 |
| 32 | 2.02 | 58.48 | 56.98 | 33.43 | - | - | - | - |
| W | 59.10 | 45.23 | 64.00 | 34.71 | - | - | - | - |
| Z | 101.54 | 41.65 | 88.16 | 10.75 | - | - | - | - |
| unplaced | 270.08 | 50.43 | 38.41 | 56.98 | 152.74 | | 44.80 | 51.41 |

Table S3: Overview of genome assemblies for all bird species included in this study. All gene annotations from this study are available at DOI: 10.5281/zenodo.12643816.

| **Species** | **Latin** | **Assembly** | **Assembly name** | **Annotation version** | **Sex** | **Total sequence length** |
|---|---|---|---|---|---|---|
| Chicken | *Gallus gallus* | https://www.ncbi.nlm.nih.gov/assembly/GCF_016699485.2/ | bGalGal1 | Ensembl | Female | 1,053,332,251 |
| Zebra finch | *Taeniopygia guttata* | https://www.ncbi.nlm.nih.gov/assembly/GCA_003957565.4 | bTaeGut1.4.pri | Refseq | Male | 1,056,271,262 |
| Downy woodpecker | *Dryobates pubescens* | https://www.ncbi.nlm.nih.gov/assembly/GCA_014839835.1 | bDryPub1.pri | This study | Female | 1,187,956,536 |
| Northern Carmine bee-eater | *Merops nubicus* | https://www.ncbi.nlm.nih.gov/assembly/GCA_009819595.1 | bMerNub1.pri | This study | Female | 1,149,356,202 |
| barn owl | *Tyto alba* | https://www.ncbi.nlm.nih.gov/assembly/GCF_018691265.1#/st | GCF_018691265.1_T.alba_DEE_v4.0 | This study | Male | 1,249,867,532 |
| Snowy owl | *Bubo scandiacus* | NA | bBubSca1 | This study | Female | 1,604,411,253 |
| Northern goshawk | *Accipiter gentilis* | https://www.ncbi.nlm.nih.gov/assembly/GCA_929443795.2 | GCF_929443795.1_bAccGen1.1 | This study | Female | 1,398,030,142 |
| California condor | *Gymnogyps californianus* | https://www.ncbi.nlm.nih.gov/assembly/GCF_018139145.2 | GCF_018139145.2_ASM1813914v2 | Refseq | Female | 1,240,248,578 |
